# Synergistic Notch–WNT Activation Underlies Pediatric Dilated Cardiomyopathy

**DOI:** 10.1101/2025.06.07.658396

**Authors:** Obed O. Nyarko, Ethan Rausch, Jared R.H. Goff, Anis Karimpour-Fard, Caitlyn S. Conard, Laura Hernandez-Lagunas, McKenna P.A. Burns, Michael R. Bristow, Joseph C. Cleveland, Matthew L. Stone, Brisa Peña, Shelley D. Miyamoto, Brian L. Stauffer, Carmen C. Sucharov

## Abstract

**Background:** Therapies for pediatric idiopathic dilated cardiomyopathy (iDCM) are extrapolated from adult heart failure despite limited efficacy, suggesting fundamental biological differences. Our prior transcriptomic studies indicate activation of developmental signaling pathways, including Notch and WNT, in pediatric iDCM; however, their mechanistic contribution remains unknown. We tested whether reactivation of Notch and WNT/β-catenin signaling drives pathological remodeling in postnatal hearts and whether pathway inhibition improves cardiac function.

**Methods:** We developed a juvenile rat model to reproduce age-dependent molecular features of pediatric iDCM using β-adrenergic stimulation (isoproterenol, ISO) and secreted frizzled-related protein-1 (sFRP1), a circulating WNT modulator elevated in children with DCM. Cardiac function was assessed by echocardiography; pathway activation by immunoblotting and transcriptomics; myocardial stiffness by atomic force microscopy. Findings were compared with explanted pediatric and adult human myocardium.

**Results:** Explanted pediatric, but not adult iDCM hearts exhibited increased nuclear and cytoplasmic Notch intracellular domain (NICD) and β-catenin. Combined ISO and sFRP1 treatment recapitulated key features of pediatric disease, including ventricular dilation, reduced ejection fraction, reactivation of the fetal gene program, and increased myocardial stiffness in the absence of fibrosis or hypertrophy. Bulk and single-nucleus RNA sequencing identified cardiomyocyte-specific activation of Notch and WNT pathways and reduced intercellular signaling diversity. Mechanistically, β-catenin silencing attenuated Notch target gene activation and pathological remodeling *in vitro*. Pharmacologic Notch inhibition reduced NICD and β-catenin accumulation, improved ventricular function, and normalized myocardial stiffness *in vivo*.

**Conclusion:** Pediatric iDCM is characterized by pathological co-activation of developmental Notch–WNT signaling pathways that are not observed in adult disease. Reactivation of this axis promotes maladaptive remodeling and myocardial stiffening, and its inhibition improves cardiac function. These findings establish developmental signaling reactivation as a central mechanism of pediatric iDCM and support age-specific therapeutic strategies.

## INTRODUCTION

Idiopathic dilated cardiomyopathy (iDCM) is one of the main causes of cardiac-related mortality in children^1^. Unfortunately, to date, there is limited evidence to support the treatment of iDCM in children, and current therapies are largely based on extrapolation of guideline-directed medical therapy (GDMT) for adult patients without proven efficacy in children^2^.

While adult DCM is predominantly characterized by chronic metabolic impairment, mitochondrial dysfunction, and fibrotic signaling pathways such as TGF-β, it lacks the enrichment of developmental signaling axes seen in children^3–5^. In stark contrast, transcriptomic profiling of pediatric DCM hearts highlights the dysregulation of WNT/β-catenin and Notch signaling pathways ^6^, which are critical for cardiogenesis but typically quiescent in the healthy postnatal heart ^7,8^.

We previously showed that pediatric iDCM secretome induces pathologic gene expression and cellular stiffness in cardiomyocytes, with elevated secreted frizzled protein 1 (sFRP1) in patient sera and hearts driving cellular stiffness in primary cardiomyocytes^9^. sFRP1 is modulates the WNT signaling pathway^10^, which regulates cellular activities through the activation of β-catenin-dependent (canonical) and independent (non-canonical) pathways^10^. WNT signaling influences Notch signaling activity in developmental contexts^7,8^. Both pathways are critical for cardiogenesis but quiescent postnatally^11,12^, yet reactivated after injury^13^.

To investigate the mechanism by which these developmental pathways drive the pediatric phenotype, we developed a novel *in vivo* model that recapitulates the specific physiological stressors found in these patients: elevated catecholamines and increased expression of sFRP1. By combining β-adrenergic stimulation (via isoproterenol) with sFRP1, we successfully modeled the synergistic co-activation of Notch and WNT signaling, thereby reproducing the unique pathological stiffness and dilation observed in the pediatric disease state. Importantly, inhibiting Notch signaling prevents pathological remodeling, cardiac dysfunction and stiffness.

## METHODS

(Detailed methods are in the Supplemental Material)

### Human Samples

Left ventricular tissue from pediatric iDCM patients undergoing transplantation and age-matched non-failing donor controls was obtained under Institutional Review Board approval at the University of Colorado in accordance with the Declaration of Helsinki. Tissue was snap-frozen for molecular analyses

### Neonatal Rat Model

Neonatal Sprague–Dawley rats were used to model pediatric cardiac remodeling. Animals were randomized to receive vehicle control, ISO, recombinant sFRP1, or combined ISO+sFRP1 treatment for 14 days. All procedures were approved by the Institutional Animal Care and Use Committee. Cardiac function was assessed by transthoracic echocardiography. Hearts were harvested for molecular, transcriptomic, and mechanical analyses.

### Molecular and Histological Analyses

Gene expression was quantified by RT-qPCR. Protein abundance and pathway activation were evaluated by immunoblotting. Histological staining was performed to assess myocardial structure and fibrosis.

### Myocardial Stiffness

Myocardial stiffness was measured using AFM. Force–indentation curves were analyzed to calculate Young’s modulus as an index of stiffness.

### Transcriptomic Analysis

Bulk RNA sequencing was performed on LV tissue. DEG and pathway enrichment analyses were conducted using established bioinformatic workflows. Single-nucleus RNA sequencing (snRNA-seq) was performed to define cell-type-specific transcriptional changes.

### Statistical Analysis

Data are presented as mean ± SEM. Comparisons between two groups were performed using unpaired Student’s t-test. Multiple group comparisons were analyzed using one-way ANOVA with appropriate post hoc testing. P<0.05 was considered statistically significant.

### Data Availability

Sequencing data will be deposited in a public repository (accession number provided upon publication). Additional data are available from the corresponding author upon reasonable request.

## RESULTS

### Human Samples

Detailed demographics for all patients included in the study are depicted in Supplemental Table 1. Median age for pediatric iDCM patients was 3.9 years (interquartile range-IQR=6.2), with an average ejection fraction (EF) of 17.33%, 58.3% were female, 8.3% were on beta-blockers, all were on angiotensin converting enzyme inhibitors/Angiotensin receptor blockers (ACEi/ARBs) and 75% were on a phosphodiesterase 3-inhibitor (PDE3i). Pediatric non-failing control (NF) subjects had a median age of 6.9 years (IQR=9.5), 28.6% were female and 14.3% were on beta-blockers. Median age for adult iDCM was 54 years (IQR=15), with an average EF of 11.58%, 50% were female, 50% were on ACEi/ARBs, 50% were on digoxin, 75% were on non-PDEi inotropes, 75% were on diuretics and 25% on PDE3i. Adult NF control subjects had a median age of 55.2 years (IQR=19.2), 50% were female, 50% were on diuretics and 25% were on beta-blockers.

### Notch and WNT signaling are activated in pediatric but not adult iDCM explanted hearts

To validate the predicted increase in Notch and WNT signaling^3^, we investigated levels of Notch intracellular domain (NICD), the constitutively active form of Notch and WNT/β-catenin protein levels in pediatric and adult iDCM hearts. We found increased levels of cytoplasmic (p=0.0013) and nuclear (p=0.0143) NICD in pediatric iDCM explanted hearts (Fig.1a,b) but not in the cytoplasmic or nuclear fractions of adult iDCM hearts (Fig.1c,d). Additionally, we found higher levels of β-catenin in the cytoplasmic (p=0.0013) and nuclear (p=0.0433) fractions of pediatric iDCM hearts (Fig.2a,b) but not in the cytoplasmic or nuclear fractions of adult iDCM hearts (Fig.2c,d).

**Figure 1.**
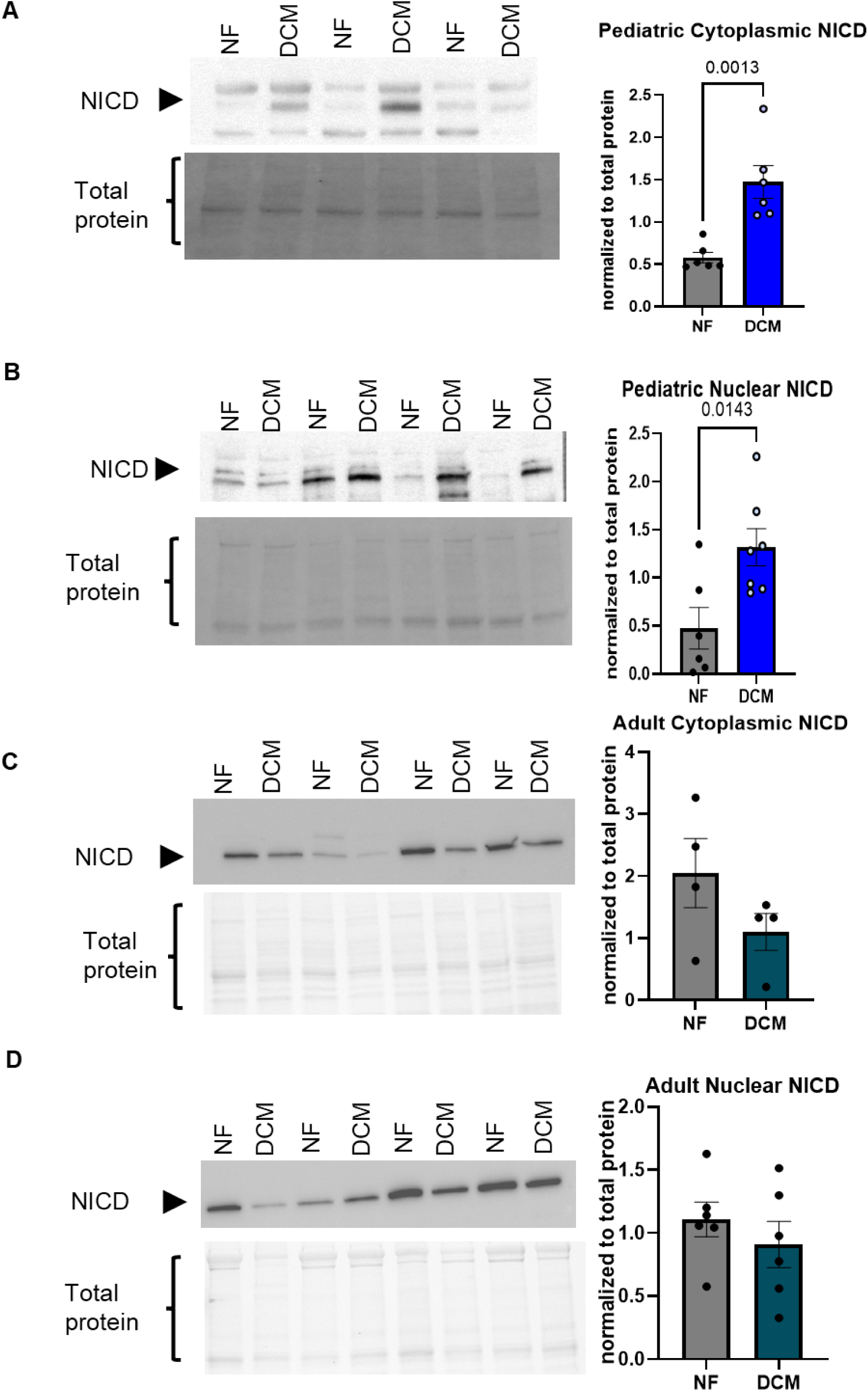
Levels of Notch are increased in pediatric iDCM hearts but do not change in adult iDCM hearts. (A-B) Immunoblot shows increased protein levels of NICD in the cytoplasm (A) and nuclear fractions (B) of pediatric NF vs iDCM human LV tissue. Quantification is shown to the right of the blots. Cytoplasmic n=6/group, nuclear n=7/group. (C-D) Immunoblot shows no changes in protein levels of NICD in adult human LV tissue. Cytoplasmic n=4/group, nuclear n=6/group. Quantification is shown to the right of the blots. Unpaired 2-tailed t test was used in analysis. Significant p-values are notated in each graph.

**Figure 2.**
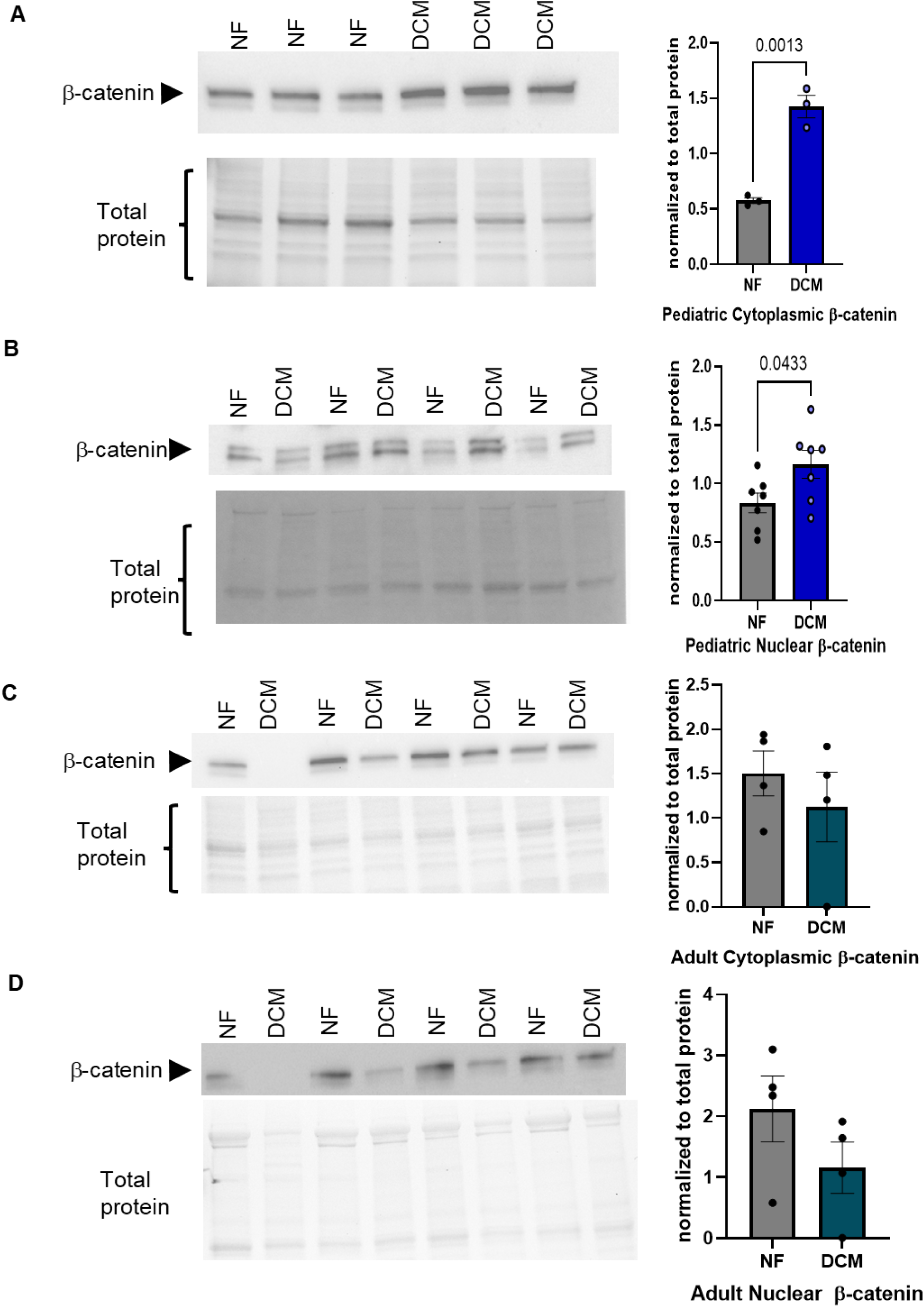
Levels of β-catenin are increased in pediatric iDCM hearts but do not change in adult iDCM hearts. (A-B) Immunoblot shows increased protein levels of β-catenin in the cytoplasm (A) and nuclear fractions (B) of NF and DCM explanted human LV tissue. Cytoplasmic n=3/group, nuclear n=7/group. Quantification is shown to the right of the blots. (C-D) Immunoblot shows no changes in protein levels of β-catenin in adult human LV tissue with quantification. N=4/group. Quantification is shown to the right of the blots. Unpaired 2-tailed t test was used in analysis. Significant p-values are notated in each graph.

### ISO+sFRP1 treatment promotes pathological remodeling *in vitro*

To recapitulate the increase in Notch and WNT signaling in the context of pediatric iDCM, we explored factors present in the secretome. The increase in circulating catecholamine can activate Notch^14,15^, whereas sFRP1 modulates WNT signaling. We first evaluated if treatment with ISO, to recapitulate an increase in circulating catecholamines, and/or sFRP1, which we previously showed increases cardiomyocyte stiffness, promoted pathological remodeling *in vitro*. Neonatal rat ventricular myocytes (NRVMs) were treated with ISO, sFRP1 or a combination of both factors for 72 hours. ISO+sFRP1 synergistically activated the fetal gene program (FGP), as measured by upregulation of *NPPA, NPPB*, and a decrease in the ratio of α-myosin heavy chain (*MYH6*) to β-myosin heavy chain (*MYH7*). *NPPA* increased in both sFRP1 (p=0.0379) and ISO+sFRP1-treated cells (p<0.0001), *NPPB* increased in ISO (p<0.0001) and ISO+sFRP1-treated cells (p<0.0001), while *MYH6/MYH7* decreased in response to sFRP1 (p<0.0001) and ISO+sFRP1 treatments (p=0.0142) (Supp.Fig.1a-c). sFRP1 expression increased in cells treated with ISO+sFRP1 (p=0.0111) (Supp.Fig.1d). Furthermore, the increase in sFRP1 gene expression in ISO+sFRP1-treated cells positively correlated with *NPPA* (p=0.0004) and *NPPB* expression (p<0.0001) and negatively correlated with *MYH6/MYH7* (p<0.0001) (Supp.Fig.2). These results suggest that a combination of ISO and sFRP1 promotes pathological remodeling *in vitro*.

### ISO+sFRP1 treatment promotes cardiac dysfunction and dilation in a young rodent model

To further validate the translational relevance of the *in vitro* model, we tested the effect of ISO+sFRP1 *in vivo*. Rats were injected with ISO and/or sFRP1 intraperitoneally for 2 weeks and then evaluated for cardiac function, (Fig.3a). Rats injected with ISO+sFRP1 had significantly reduced EF (p<0.0001) when compared to vehicle-treated control rats, as well as when compared to rats injected solely with ISO or sFRP1 (Supp.Table 2). Left ventricular volume in systole (LV vol(s)) increased in ISO+sFRP1-treated rats compared to vehicle-treated controls (p=0.0002), ISO-treated rats or sFRP1-treated rats. Similarly, left ventricular internal diameter in systole (LVID(s)) increased in ISO+sFRP1 rats compared to vehicle-treated controls (p<0.0001), ISO-treated rats or sFRP1-treated rats (Supp.Table 2). There were no significant differences in left ventricular anterior and posterior wall thickness for any of the treated groups compared to controls (Supp.Table 2). Diastolic parameters, LV vol(d) and LVID(d), followed the same trend as systolic parameters (Supp.Table 2). Morphometric data showed a significant increase in whole heart to body weight or tibia length and LV weight to body weight or tibia length in ISO+sFRP1 rats compared to vehicle-treated controls (Supp.Fig.3a, b). Functional alterations were not restricted to male animals; changes were also observed in females (Supp.Fig.3c).

**Figure 3.**
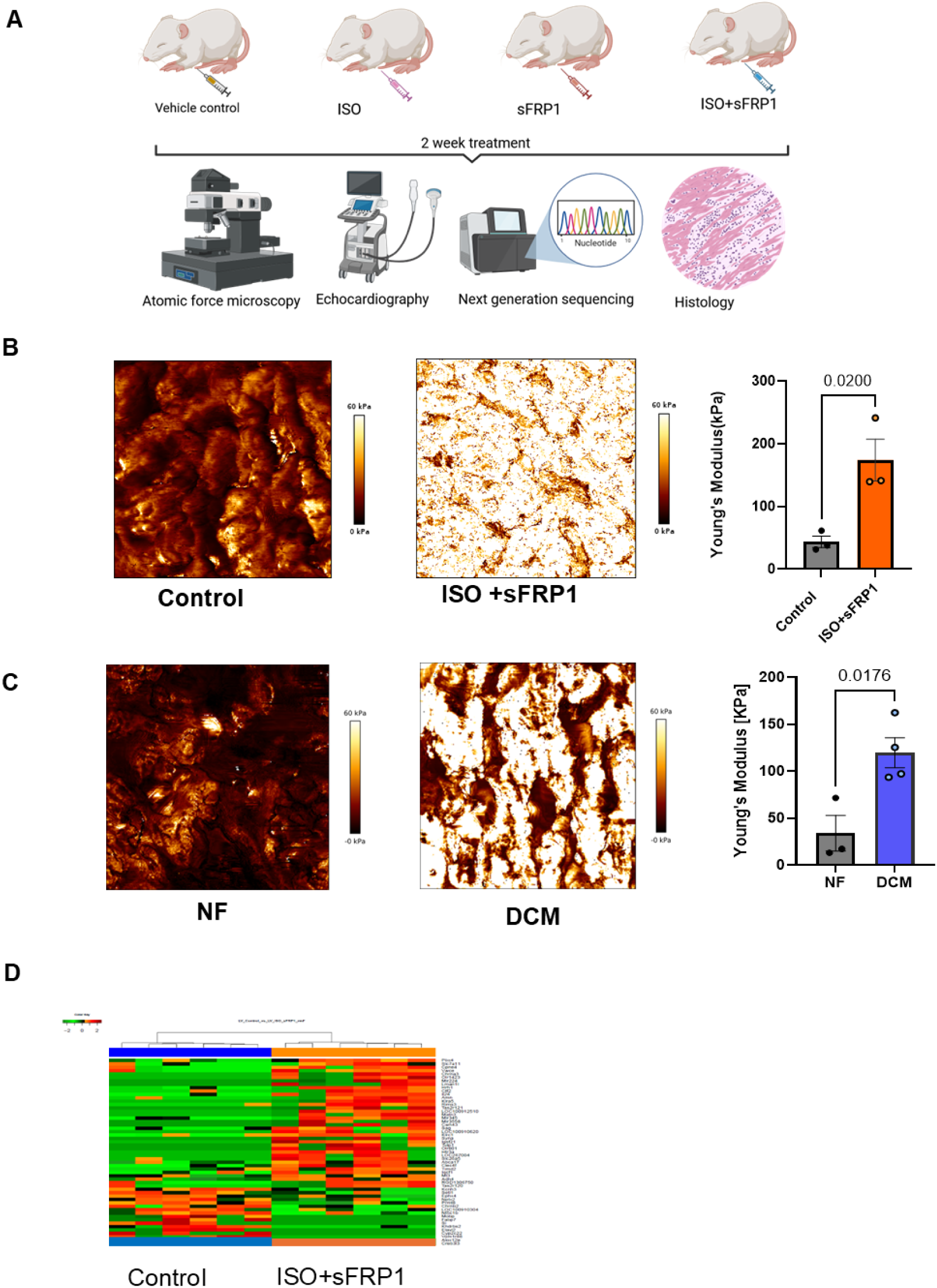
Treatment of ISO+sFRP1 *in vivo* recapitulates the increased stiffness observed in human pediatric iDCM hearts. (A) Schematic of the experimental plan for intraperitoneal injections of rats treated with vehicle control, ISO, sFRP1 and ISO+sFRP1. (B) Elasticity/stiffness (Young’s modulus) in rats treated with ISO+sFRP1 compared to controls. Quantification is shown to the right. *n* =3 per group. Unpaired 2-tailed t test was used in analysis. (C) Elasticity/stiffness (Young’s modulus) in explanted pediatric iDCM hearts compared to non-failing controls. Quantification is shown to the right. NF n=3, DCM n*=*4. Unpaired 2-tailed t test was used in analysis. (D) Heatmap representing unsupervised hierarchical clustering of the top 50 significantly differentially expressed genes (DEGs) in bulk RNA sequencing. *n*=6/group; Welch’s *t*-test *p*<0.05.

As in the *in vitro* expression studies, *NPPA* expression increased in sFRP1- (p=004) and ISO+sFRP1-treated rats (p=0.0035) compared to vehicle-treated controls (Supp.Fig.4a). However, *NPPB* expression only increased in sFRP1-treated rats (p=0.0011) and not in response to ISO+sFRP1 treatment (Supp.Fig.4b). *MYH6/MYH7* ratio decreased in both sFRP1 (p=0.0272) and ISO+sFRP1 rats (p=0.049) (Supp.Fig.4c). A simple linear regression was performed to evaluate the relationship between pathological cardiac remodeling, evidenced by reactivation of the FGP, and ejection fraction. We found that the relative expression of *NPPA*, but not *NPPB*, was significantly correlated with a decrease in EF (p=0.0039) (Supp.Fig.5a,b), whereas the ratio of *MYH6/MYH7* trended towards a positive correlation (higher *MYH6/MYH7* correlated with higher EF) (p=0.0825) (Supp.Fig.5c). These results, together with the *in vitro* data, provide evidence that ISO+sFRP1 promotes pathological remodeling *in vitro* and cardiac dysfunction associated with left ventricular dilation *in vivo*.

To investigate the translational significance of the rat ISO+sFRP1 DCM model and endogenous expression pattern of sFRP1, we investigated cardiac mRNA and serum sFRP1 levels in rats relative to our published data in human pediatric DCM patients^6,9^. We observed a significant increase in cardiac sFRP1 mRNA levels in ISO+sFRP1 rats (p=0.0003) (Supp.Fig.6a) that was negatively correlated with EF (p=0.0121) (Supp.Fig.6b). Moreover, levels of sFRP1 in the serum of ISO+sFRP1 rats were significantly higher than vehicle-treated controls (Supp.Fig.6c) (p=0.0146). There was a trend to negative correlation when analyzing levels of serum circulating sFRP1 and EF (p=0.0888) in rats (Supp Fig.6d). The observed increase in rat cardiac and serum sFRP1 levels was comparable to our previously published human data^9^.

### ISO+sFRP1 promotes increased stiffness in the absence of myocyte hypertrophy or fibrosis

We previously showed that DCM serum and sFRP1 treatment promote stiffness in NRVMs assessed by atomic force microscopy (AFM)^9^. To further explore the role of sFRP1 in promoting stiffness *in vivo*, we investigated myocardial stiffness in our model. Critically, ISO+sFRP1 promoted a 2-fold increase in LV stiffness when compared to vehicle-treated control rats (p=0.02) (Fig.3B), matching the stiffness observed in explanted pediatric DCM hearts (Fig.3C), despite absent fibrosis or hypertrophy (Supp.Fig.8). This uncoupling of stiffness from fibrosis suggests cytoskeletal or ECM proteoglycan remodeling as a pediatric-specific mechanism. Stiffness is specific to ISO+sFRP1 treatment as changes in myocardial stiffness were not observed in rats treated with ISO or sFRP1 (Supp.Fig.7a-d).

We and others previously showed that pediatric iDCM patients have minimal fibrosis and lack myocyte hypertrophy^16,17^. Compared to vehicle-treated controls, ISO+sFRP1 rats did not exhibit increased fibrosis, as determined by Masson’s trichrome staining (Supp.Fig.8a). Additionally, RT-qPCR analysis of the rat LV tissue demonstrated no significant upregulation of profibrotic genes, including *COL1A1, COL3A1, Galectin-3, TGFB1*, *TGFB2* and *TIMP1-4* (Supp.Fig.8b). Furthermore, myocyte hypertrophy was not observed in ISO+sFRP1 rats compared to controls (Supp.Fig.8c).

### Notch and WNT signaling are altered in ISO+sFRP1 rats and in pediatric DCM hearts

To better understand the pathophysiology of ISO+sFRP1-mediated dysfunction and identify the mechanisms underlying the disease, we performed bulk RNA sequencing analysis of the LV of rats treated with ISO and/or sFRP1 compared to vehicle-treated controls (n=6/group). Of the 14,350 genes identified, there were 286 differentially expressed genes (DEGs) (p<0.05) in ISO+sFRP1 rats compared to vehicle-treated controls (113 up-regulated and 173 down-regulated) (Supp.Table 3). Unsupervised hierarchical clustering separated control and ISO+sFRP1 treatment groups based on the gene expression profile. Unsupervised hierarchical cluster heatmaps of the top 50 DEGs in ISO+sFRP1- (Fig.3d), sFRP1- or ISO-treated rats (Supp.Fig.9a,b) compared to vehicle-treated controls were generated and showed a clear separation between individual groups and control. Gene set enrichment analysis (GSEA) pathway analysis identified unique patterns of pathway activation in ISO+sFRP1 compared to control such as microtubule cytoskeleton organization (Fig.4a).

**Figure 4.**
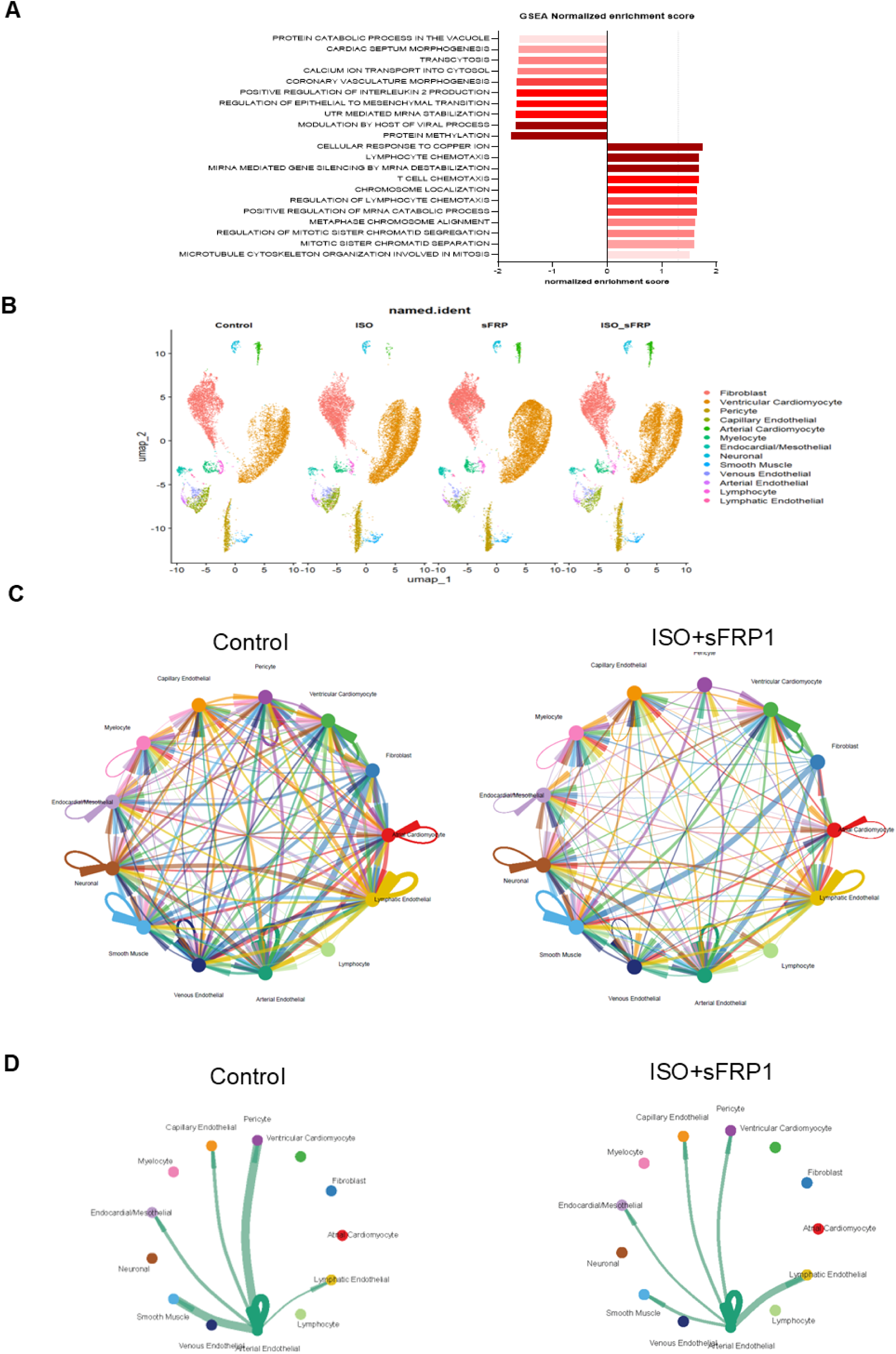
Next generation sequencing analysis highlighting Notch signaling and global reduction in signaling diversity in the ISO+sFRP1 model. (A) Gene set enrichment analysis of bulk RNA-sequencing of rats treated with ISO+sFRP1 compared to vehicle-treated controls. (B) Representative UMAP plot from single-nuclei RNA sequencing of LV tissue from Controls, ISO-, sFRP1- or ISO+sFRP1-treated rats, 3 rats pooled per group; 1 library per condition. A total of 9214 nuclei in control, 11588 nuclei in ISO-, 13758 nuclei in sFRP1-, and 11127 in ISO+sFRP1-treated rats were identified after cells were pooled. 26 distinct cell clusters were identified including 8 CMs and 5 FBs. (C) CellChat analysis of the total number of cellular interactions in ISO+sFRP1 treated rats compared to controls. (D) Cellchart intercellular communication of the NOTCH signaling pathway network of rats treated with vehicle control, ISO, sFRP1 and ISO+sFRP1. UMAP, uniform manifold approximation and projection; CM, cardiomyocytes; FB, fibroblast; NK, Natural Killer cells; B/T, B cells and T cells;

To investigate commonalities in hearts from children with DCM and our rat model, we assessed similarities between DEGs from ISO+sFRP1 rats, pediatric DCM hearts, and from our previously published data on pediatric DCM serum-treated NRVMs^9^. Transcriptomic analysis revealed a substantial overlap in DEGs in the LV tissue of ISO+sFRP1-treated rats and human DCM hearts or serum-treated cells. Notably, 96% of DEGs in the LV tissue of ISO+sFRP1-treated rats were also observed in the human DCM heart (65% unique only to human LV DEGs, 31% shared between human LV and rat LV DEGs with only 4% DEGs unique to rats) (Supp.Fig.10a). Additionally, 43% of DEGs were common between the LV tissue of ISO+sFRP1-treated rats and NRVMs treated with human DCM serum (57% unique to NRVMs treated with human DCM serum DEGs, 1% shared between NRVMs treated with human DCM serum DEGs and rat LV DEGs with 42% DEGs unique to rats) (Supp.Fig.10b). Notch signaling was one of the top dysregulated pathways associated with commonly upregulated genes in DCM serum-treated NRVMs and LV tissue of ISO+sFRP1-treated rats (Supp.Fig.10c).

### Single-nuclei RNA sequencing (snRNA-seq) identifies activation of Notch and β-catenin signaling in response to ISO+sFPR1

To further characterize the cell types and biological mechanisms altered at the individual cellular level, we performed single-nuclei RNA sequencing of LV tissue from ISO and/or sFRP1, and vehicle-treated control rats (n=3 rats/condition pooled to generate n=1 library/group). A total of 9,214 nuclei in control, 11,588 nuclei in ISO-, 13,758 nuclei in sFRP1-, and 11,127 in ISO+sFRP1-treated rats were identified after cells were pooled. Clusters were annotated based on canonical cell marker expression (Supp.Table 4.1). We assessed the distribution of cell-type annotated nuclei across all samples and identified 26 transcriptionally distinct cell-type clusters marked by canonical gene markers (Fig.4b)^18,19^. A sub-analysis of ventricular cardiomyocytes identified eight different sub-clusters (Supp.Fig.11a). Within this dataset, ISO+sFRP1 was associated with an expansion of ventricular cardiomyocyte subcluster 2 (the cluster with the most relative proportion of ventricular cardiomyocytes) and enrichment of Notch signaling pathway with predicted *CTNNB1* (β-catenin) transcriptional control (Supp.Fig11b,c, Supp.Table 4.2), while subcluster 6 (cluster with the 2^nd^ most relative proportion of ventricular cardiomyocytes) exhibited extracellular matrix (ECM) proteoglycan enrichment with *Notch1* as a predicted regulator (Supp.Fig.11d, e, Supp.Table 4.3). Pathway analysis was also performed on the DEGs of the stromal cell types such as fibroblast, pericytes and endothelial cells which further predicted alterations in ECM proteoglycans but no alterations in Notch/WNT signaling in these cell types (Supp.Fig. 12).

### Global Reduction in Signaling Diversity in the ISO+sFRP1 Model

Analysis of the total number of cellular interactions using CellChat suggest a reduction in inferred intercellular communication in the ISO+sFRP1 model compared to vehicle-treated controls (Fig.4c and quantified in Supp.Fig.13a). Visualization of the Notch signaling network revealed a marked decrease in network density and complexity, with fewer active ligand-receptor connections between most cell types in the ISO+sFRP1 treated rats compared to controls (Fig.4d).

### Notch and WNT signaling are activated in ISO+sFRP1 *in vitro* and *in vivo* treated neonatal rats and NRVMs

To test if ISO+sFRP1 promotes activation of Notch and WNT signaling, we investigated the protein levels of NICD and β-catenin, and expression of genes in these pathways. NICD protein was increased in cytoplasmic (p=0.0053) (Fig.5a) and nuclear fractions (p=0.0086) of ISO+sFRP1 rat hearts (Fig.5b), and in cytoplasmic (p=0.0371) (Fig.5c) and nuclear (p =0.0119) (Fig.5d) fractions of ISO+sFRP1-treated NRVMs. In parallel, we also observed increased expression of the Notch-target genes in NRVMs, and cardiac tissue of rats treated with ISO+sFRP1, which recapitulated pediatric DCM heart data. Specifically, *Notch1* (p=0.0398) and *HEY1* (p=0.0467) were up regulated in ISO+sFRP1 rat hearts (Supp.Fig.14a); *Notch1* (p=0.0276) together with the expression of Notch target genes *HEY1* (p=0.0006), *HEY2* (p=0.0124), *HES1* (p=0.0361) and *HEYL* (p=0.0436) were increased in NRVMs treated with ISO+sFRP1 compared to vehicle-treated controls (Supp.Fig.14b). *Hey1*, was upregulated in pediatric DCM hearts (1.55x, q=0.02 from RNA-seq data)^6^. There was no increase in expression of other Notch receptor genes, Notch2-4 (Supp. Fig.14c). Expression of *Notch 1* and Notch-target genes was not altered in ISO or sFRP1-treated NRVMs (Supp.Fig.15a-e).

**Figure 5.**
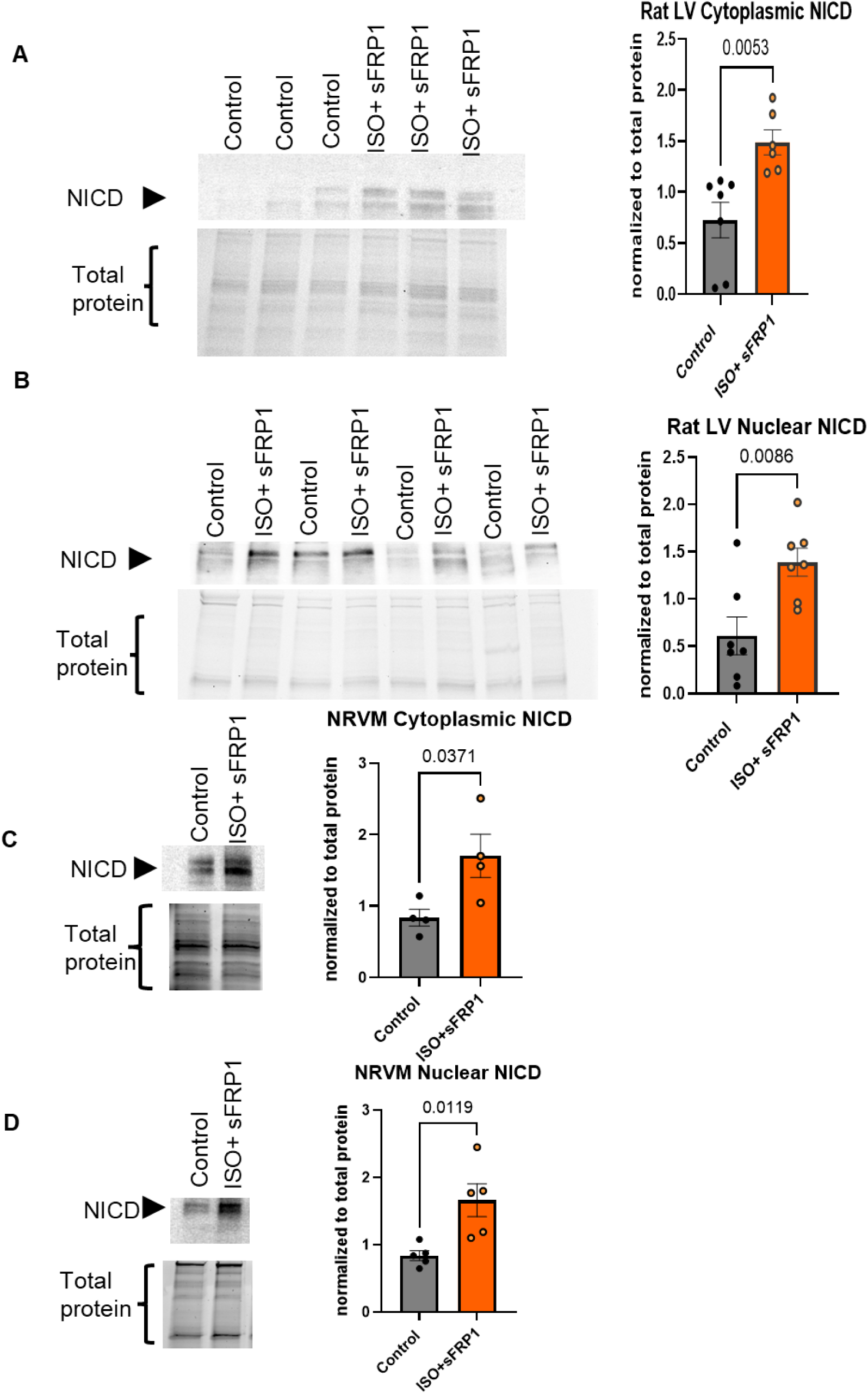
Levels of NICD are increased in nuclear and cytoplasmic fractions in *vivo* and *in vitro*. (A, B) Immunoblot shows increased protein levels of NICD in the cytoplasm (A) and nuclear fractions (B) of ISO+sFRP1 vs vehicle-treated control rat LV tissue. Cytoplasmic n=6/group, nuclear n=7/group. Quantification is shown to the right of the blots. (C, D) Immunoblot shows the increased protein levels of NICD in the cytoplasm (C) and nuclear fractions (D) of ISO+sFRP1 vs vehicle-treated NRVMs. Cytoplasmic n = 4 independent NRVM preps, nuclear n = 5 independent NRVM preps. Quantification is shown to the right of the blots. Unpaired 2-tailed t test was used in analysis. *P*-values are notated in each graph.

sFRP1 is known to modulate the WNT/β-catenin pathway in a context-dependent manner^20^. We found higher levels of β-catenin in the cytoplasmic (Fig.6a) (p=0.0099) and nuclear fractions (Fig.6b) (p-value=0.0191) of ISO+sFRP1 rat hearts, and in the cytoplasmic (Fig.6c) (p=0.0101) and nuclear fractions (Fig.6d) (p=0.0039) of ISO+sFRP1-treated NRVMs. Additionally, we also observed an increase in expression of *CTNNB1* (p=0.0263) in ISO+sFRP1 rats and ISO+sFRP1-treated NRVMs (p=0.0460) (Supp.Fig.16a,b). Similar to what we observed when evaluating Notch signaling, the increase in *CTNNB1* levels was unique to ISO+sFRP1-treated NRVMs and was not observed in response to ISO or sFRP1. (Supp.Fig16c).

**Figure 6.**
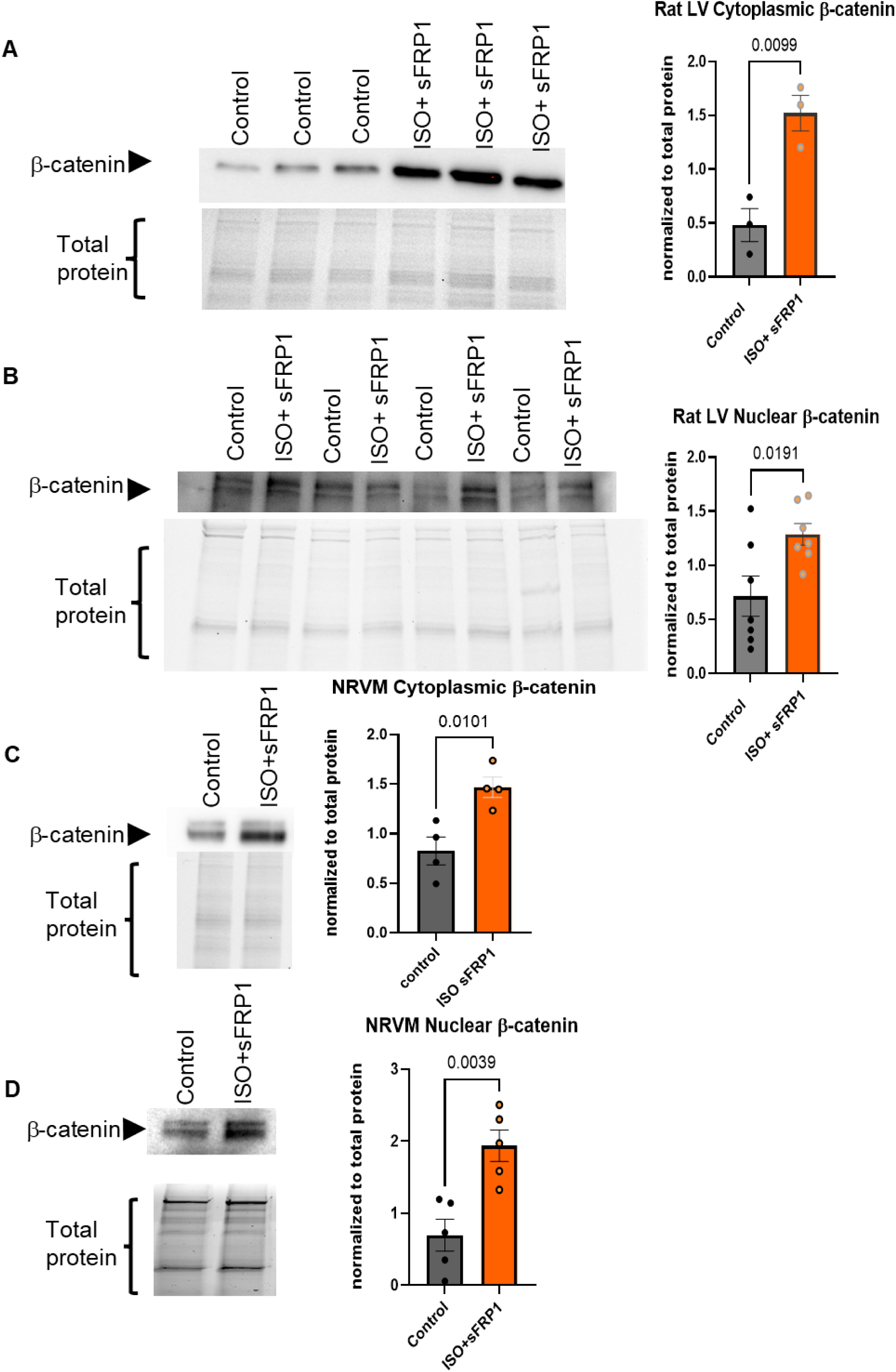
Levels of β-catenin are increased in nuclear and cytoplasmic fractions *in vivo* and *in vitro*. (A, B) Immunoblot shows increased protein levels of β-catenin in the cytoplasm (A) and nuclear fractions (B) of ISO+sFRP1 vs vehicle-treated control rat LV tissue. Cytoplasmic n=3/group, nuclear, n=7/group. Quantification is shown to the right of the blots. (C, D) Immunoblot shows increased protein levels of β-catenin in the cytoplasm (C) and nuclear fractions (D) of ISO+sFRP1 vs vehicle-treated NRVMs. Cytoplasmic and nuclear fractions from n=4 independent NRVM preps. Quantification is shown to the right of the blots. Unpaired 2-tailed t test was used in analysis. *P*-values are notated in each graph.

### β-catenin contributes to Notch activation in ISO+sFRP1-treated cells

Others have reported an interplay between Notch and β-catenin signaling^21,22^, so we next evaluated the contribution of β-catenin to Notch activation and vice versa. siRNA-mediated down-regulation of *CTNNB1* prevented ISO+sFRP1-mediated activation of Notch target gene expression by blunting expression of *HEY1* (p<0.0001), *HEY2* (p=0.0026), *HES1* (p=0.0167) and *HEYL* (p=0.0401) (Supp. Fig.17a). Additionally, down-regulation of *CTNNB1* also prevented ISO+sFRP1-mediated activation of the FGP by blunting the increase in expression of *NPPA* (p=0.0063) and *NPPB* (p=0.0003), and by increasing the ratio of *MYH6/MYH7* (p=0.0446) (Supp.Fig.17b). Down-regulation of *CTNNB1* also reduced the ISO+sFRP1-mediated increase in cytoplasmic (p=0.0041) (Fig.7a) and nuclear (p=0.0063) (Fig.7b) NICD. The effect of down-regulating CTNNB1 on β-catenin levels is shown in Fig.7c,d.

**Figure 7.**
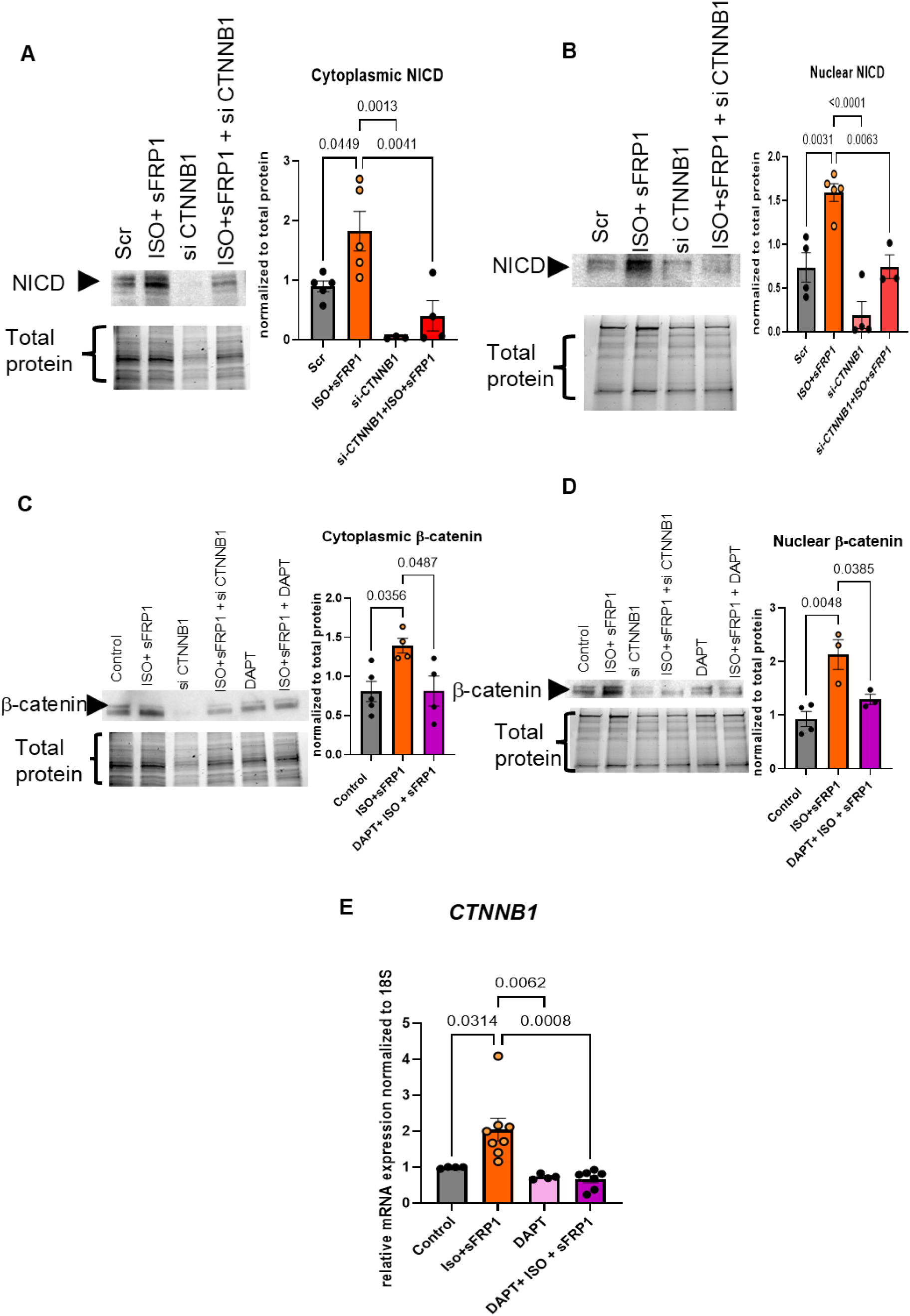
β-catenin and Notch co-activation in ISO+sFRP1-treated cells. (A, B) Immunoblot shows reduced protein levels of NICD in the cytoplasm (A) and nuclear fractions (B) of NRVMs treated with ISO+sFRP1+si-CTNNB1 compared to ISO+sFRP1+scramble control, scramble control and si-CTNNB1. n=3 independent NRVM preps/group. Quantification is shown to the right of the blots. (C-D) Immunoblot shows the reduced protein levels of β-catenin in the cytoplasm (C) and nuclear fractions (D) of ISO+sFRP1+DAPT compared to ISO+sFRP1-treated, DAPT and vehicle-treated control NRVMs. *n*=4 independent NRVM preps/group. Quantification is shown to the right of the blots. (E) RT-qPCR shows reduced CTNNB1 gene expression in ISO+sFRP1+DAPT compared to ISO+sFRP1-treated, DAPT and vehicle-treated control. Gene expression was normalized to 18S, and data are presented as a relative fold change to Controls. Tukey’s multiple comparisons test was used for all data sets. Control, n=4, ISO+sFRP1 n=8, DAPT n=4 and ISO+sFRP1+DAPT n=7 Scr, scrambled control;

### Notch antagonism prevented pathological remodeling *in vitro* and cardiac dysfunction associated with dilation *in vivo*

We also tested the effect of DAPT, a γ-secretase inhibitor known for reducing the levels of NICD, on β-catenin levels and activation of the FGP. DAPT, as expected, prevented or decreased the activation of Notch target genes *HEY1* (p<0.0001), *HEY2* (p=0.0004), *HES1* (p<0.0001) and *HEYL* (p<0.0001) by ISO+sFRP1 (Supp.Fig.17c). We next tested whether Notch inhibition affected activation of the FGP. As shown in Supp. Fig.17d. DAPT prevented the reactivation of FGP by ISO+sFRP1 by blunting the increased expression of *NPPA* (p=0.025), and *NPPB* (p=0.0014), and by increasing the ratio of *MYH6/MYH7* (p=0.007). Additionally, DAPT prevented ISO+sFRP1-mediated increase in cytoplasmic (p=0.0487) (Fig.7c) and nuclear (p=0.0385) (Fig.7d) β-catenin. DAPT prevented the increase in *CTNNB1* gene expression (p=0.008) (Fig.7e). These data support the premise that the co-activation of Notch and β-catenin promotes pathological remodeling.

Lastly, we investigated the role of DAPT in preventing cardiac dysfunction associated with ventricular dilation in our ISO+sFRP1 *in vivo* model (Fig.8a). DAPT prevented the reduction of EF by ISO+sFRP1 (p=0.0048) and trended towards inhibiting the increase in LV vol(s) (p=0.088) and LVID(s) (p=0.0935) (Fig.8b-d). Additionally, DAPT also abrogated the increase in myocardial stiffness mediated by ISO+sFRP1 (p=0.0179) (Fig.8e). This increase in myocardial stiffness negatively correlated with cardiac dysfunction (p=0.0271) (Fig.8f). The correlation between EF and stiffness suggests that Notch inhibition may improve both contractile function and myocardial properties, though whether stiffness reduction is direct (via cytoskeletal changes) or indirect (via reduced wall stress) requires further investigation

**Figure 8.**
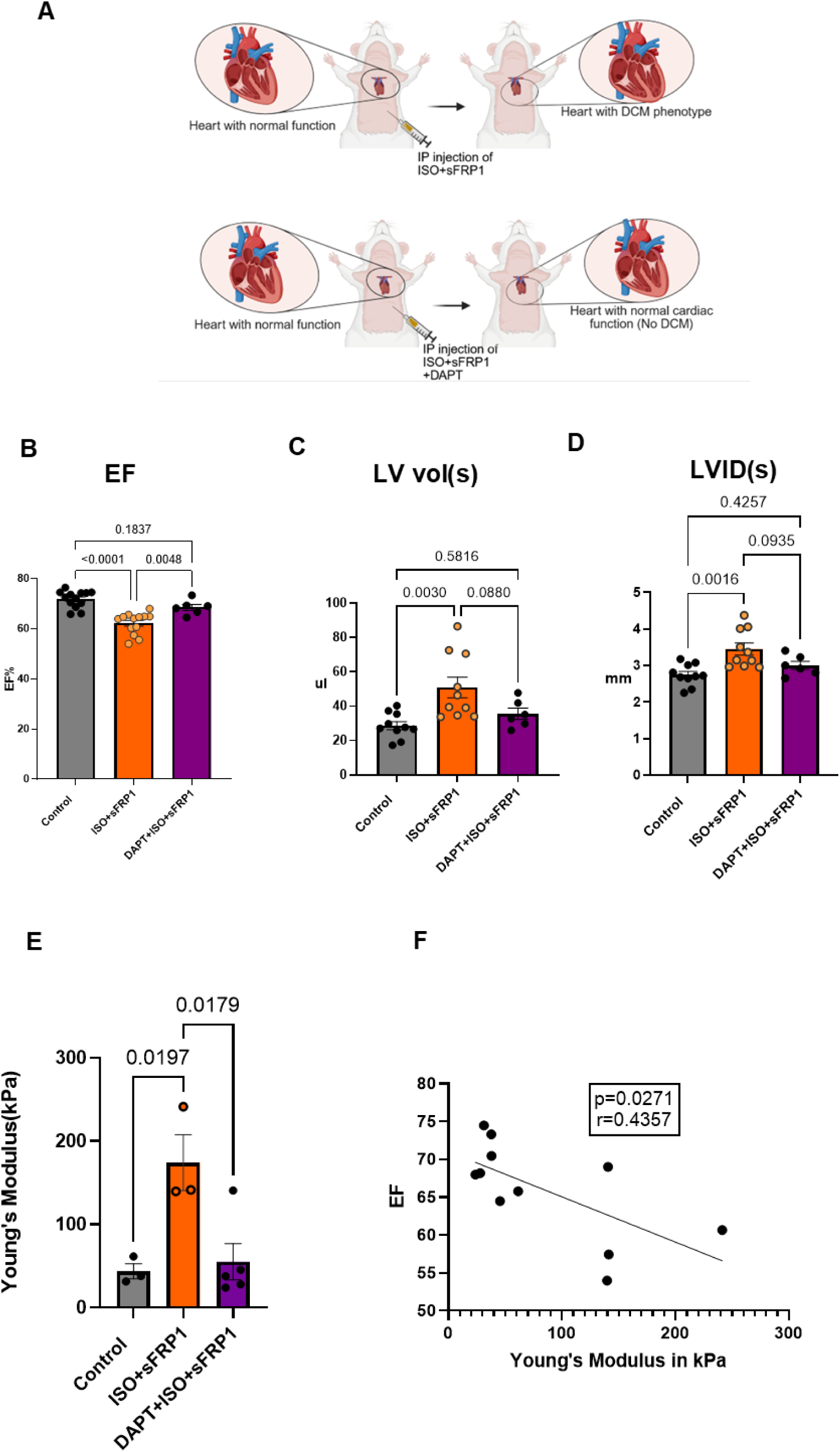
Inhibiting Notch prevents cardiac dysfunction and stiffness in ISO+sFRP1-treated rats. (A) Schematic of the experimental plan for rats treated with ISO+sFRP1 +/-DAPT showing cardiac dysfunction is prevented. (B) EF of neonatal rats treated with Control, ISO+sFRP1 and DAPT+ISO+sFRP1. (C) LV vol(s) of neonatal rats treated with Control, ISO+sFRP1 and DAPT ISO+sFRP1. (D) LVID(s) of neonatal rats treated with Control, ISO+sFRP1 and DAPT ISO+sFRP1. Control n=13, ISO+sFRP1 n=13, DAPT+ISO+sFRP1 n=6. (E) Elasticity/stiffness (Young’s modulus) in rats treated with DAPT+ISO+sFRP1 compared to ISO+sFRP1 treated rats and controls. Quantification is shown. Control n=3, ISO+sFRP1 n=3 and DAPT+ISO+sFRP1 n=4. (F) Simple linear regression of cardiac function (EF) and stiffness (Young’s modulus in kPa). Tukey’s multiple comparisons test was used for all data sets. *P*-values are notated in the graph.

## DISCUSSION

### Pediatric iDCM Exhibits a Distinct Developmental Signaling Signature

The clinical management of pediatric heart failure has relied largely on therapies derived from adult studies, despite limited efficacy in children. Our prior findings support the premise that pediatric iDCM is biologically distinct from adult disease, including minimal fibrosis/hypertrophy^6,16^, and suggest activation of Notch and WNT^6^. In the current study, analysis of explanted myocardium demonstrated robust nuclear and cytoplasmic accumulation of NICD and β-catenin in pediatric, but not adult, iDCM, identifying co-activation of Notch and WNT/β-catenin as a defining molecular feature of the pediatric failing heart. These pathways, essential during cardiogenesis but largely quiescent postnatally, appear pathologically activated in pediatric disease.

### Modeling Pediatric-Specific Pathophysiology

To test causality, we developed a juvenile rat model incorporating two circulating factors elevated in pediatric iDCM: catecholaminergic stress and sFRP1. Combined ISO+sFRP1 exposure recapitulated key hallmarks of pediatric disease, including ventricular dilation, reduced ejection fraction, fetal gene reactivation, and increased myocardial stiffness in the absence of fibrosis or hypertrophy. Importantly, this model reproduced the Notch–WNT co-activation observed in human samples, supporting a mechanistic link between developmental signaling reactivation and functional decline.

Our model has an over 10% absolute reduction in EF which is a significant and important indicator of decline and physiological change in the heart’s pumping ability. Such declines in cardiac function in humans are associated with increased risk of all-cause mortality^23^. Additionally, rodent echocardiographic data indicates that EF declines with age ^24,25^, meaning young rats typically have higher baseline EF. The relative EF reduction in ISO+sFRP1 rats likely reflects early dysfunction, not preserved contractility^24,25^. The dilated chambers are one of the primary criteria for DCM and the significant increase in dilation. Considering we observed significant changes in Echo parameters after only two weeks of treatment with ISO+sFRP1, our model is likely to represent the early onset of iDCM. The two-week treatment duration captures initiation of maladaptive remodeling, evidenced by FGP activation, LV dilation, and stiffness, before fibrotic scarring. This temporal window is clinically relevant, as pediatric iDCM often presents acutely with rapid decompensation. Chronic studies (4-8 weeks) are ongoing to assess progression to end-stage HF.

### Notch–WNT Crosstalk Drives Pathological Remodeling

The WNT/β-catenin pathway has a biphasic role in cardiac development and differentiation, promoting cardiac differentiation before gastrulation and inhibiting cardiac development after gastrulation^20^. Additionally, although stabilizing β-catenin promotes proliferation of mammalian cardiomyocytes^26^, it is also required for stress-induced cardiac hypertrophy^27^. These studies highlight the complexity of WNT/β-catenin signaling in the heart. In our study, we show that WNT/β-catenin contributes to a pathological phenotype, suggesting the activation of this pathway in post-natal hearts is associated with maladaptive remodeling and cardiac dysfunction in pediatric but not adult iDCM.

Notch signaling is fundamentally involved in cell-cell communication and has been implicated in cardiac development, function and cardiac birth defects^28^. Notch target genes contribute to cardiac development and disease^29^, and aberrations in the Notch signaling pathway are associated with congenital heart defects^11^. In addition, Notch signaling is activated after cardiac injury^30^, and promotes cardiac regeneration^31^, and cellular stiffness^32^. These suggest that Notch might be beneficial in regenerative, compliant environments such as the developing heart, but potentially detrimental in stiff, non-regenerative conditions such as the postnatal heart.

To understand how this signaling drives pathology at the tissue level, we utilized single-nuclei RNA sequencing (snRNA-seq) and CellChat analysis. These revealed a "signaling paradox" in the pediatric failing heart. Globally, the ISO+sFRP1 hearts exhibited a breakdown of homeostatic communication, evidenced by a marked reduction in the predicted total number of intercellular interactions as the heart transitions to a failing phenotype. While the healthy heart maintains a high volume of diverse homeostatic signals, the diseased state is characterized by a marked reduction of cardiac networking, where fewer total pathways that are active across the myocardial and non-myocardial cell populations^33^. Similar communication breakdown occurs in hypertrophic cardiomyopathy, where altered ECM signaling disrupts homeostasis^34^.

Interestingly, our bulk and single-nuclei transcriptomics demonstrated a significant upregulation of intracellular *Notch1* mRNA and increased NICD protein in the cytoplasmic and nuclear fractions of ISO+sFRP1 hearts with decreased interaction frequency and relative strength. Despite reduced intercellular communication, intracellular Notch1 mRNA and NICD protein were elevated, suggesting ligand-independent Notch activation that may be driven by catecholamine stress^35^.

In the heart, Notch and WNT pathways are tightly regulated during embryogenesis, where they coordinate cardiac progenitor specification and morphogenesis^31^. However, in diseased states, their reactivation often contributes to maladaptive remodeling^36^. Previous studies have shown that WNT/β-catenin signaling can enhance Notch pathway activation through direct transcriptional regulation of Notch ligands^21,22^ while Notch signaling has been reported to modulate β-catenin stability and nuclear translocation^37^. This bidirectional crosstalk may underline the robust transcriptional and protein response we observed in our data. Interestingly, in our model, we detected an expansion of cardiomyocyte cluster 2 in ISO+sFRP1 hearts. In this cluster, there is an upregulation of Notch signaling, and *CTNNB1* is predicted to be one of the top transcriptional regulators, supporting our findings that ISO+sFRP1 treatment results in WNT/Notch activation. Thus, the expansion of ventricular cardiomyocyte nuclei in cluster 2 may reflect both a compensatory survival mechanism and a shift toward a transcriptionally dysregulated state driven by concordant Notch-WNT pathway synergy. In congruence, ventricular cardiomyocyte cluster 6 projects *Notch1* as one of the top transcription factors predicted to control expression of upregulated genes. This suggests that one cluster may be more β-catenin-regulated and the other, Notch-regulated.

Our observation that both pathways are upregulated in ISO+sFRP1-treated cells, rats, and human DCM hearts suggests a cooperative role in driving pathological gene expression programs associated with cardiac dysfunction and ventricular dilation *in vivo*. These findings support a model in which combined β-adrenergic stimulation and increased WNT signaling activate Notch–WNT signaling, contributing to sustained pathological remodeling and cardiac dysfunction.

### sFRP1 regulation of WNT signaling

sFRP1’s WNT-agonist effect in our model aligns with myocardial infarction studies^38^, ^39^where prolonged exposure and/or increased concentration of sFRP1 can promote β-catenin stabilization and WNT signaling activation^39^. Our results agree with these findings, as initial exposure (3 hours to 24 hours) to ISO+sFRP1 did not increase β-catenin levels (not shown), but persistent (72 hours) exposure increased levels of β-catenin in cardiomyocytes. The mechanisms associated with sFRP1-mediated stabilization of β-catenin are unclear; sFRP1 may directly form complexes with frizzled receptors, possibly mimicking the effects of WNT ligands or displacing inhibitory interactions, thus activating downstream signaling^40^. Furthermore, it has been proposed that the duration or levels of exposure to sFRP1 may stabilize certain WNT ligands or alter receptor dynamics to enhance β-catenin accumulation^39^.

Our results also suggest a context-dependent function of sFRP1, in which sFRP1 may differentially modulate WNT signaling under adrenergic stress. In cancer, chronic adrenergic stress has been shown to promote sFRP1 secretion, enhancing tumor growth through β-catenin activation^39^. Hence targeting sFRP1 has been explored as a potential therapeutic target for patients with adenocarcinoma^41^. Our findings extend these observations by demonstrating that simultaneous β-adrenergic stimulation and WNT pathway modulation can synergistically trigger maladaptive gene expression patterns and cardiac dysfunction. In this context, ISO appears to act as a modulator of the response to sFRP1 and may modify the cellular context, allowing WNT signaling components to become activated despite the presence of a known antagonist, perhaps through non-canonical or compensatory mechanisms.

### Notch as a potential therapeutic target in pediatric iDCM

Although Notch signaling is essential for homeostasis and unlikely to be broadly suppressed therapeutically^42^, our data implicate downstream effectors, particularly cytoskeletal remodeling and stiffness, as targetable. Our snRNA-seq data implicate ECM proteoglycans (enriched in ventricular CM cluster 6) as candidate mediators. Future studies using proteomics and AFM on isolated cardiomyocytes will dissect cytoskeletal vs. ECM contributions and can investigate whether Notch-mediated ECM proteoglycan expression or titin modifications drive pediatric DCM stiffness.

Additionally, our *in vivo* model recapitulates other aspects of pediatric iDCM, such as no difference in cardiac *NPPB* levels, minimal fibrosis or lack of myocyte hypertrophy^6,16^. While fibrosis was not observed in our model, we detected an increase in myocardial stiffness, which recapitulates our prior *in vitro* studies in cells treated with sFRP1^9^. The increase in stiffness in the absence of fibrosis highlights that these two processes can be independently regulated. Notably, both WNT and Notch signaling have been implicated in the regulation of ECM dynamics and fibroblast activation in adult cardiac fibrosis^43^. Although ECM remodeling and myocardial stiffness have been detected in the hearts of children with iDCM^5,26^, it is possible that modifications of sarcomeric proteins, titin isoform expressions or modifications might have an impact on stiffness which deserves further investigations.

Taken together, our findings suggest myocardial stiffness as an important contributor to pediatric iDCM and support WNT and Notch signaling pathways as important drivers of the disease in children.

## CONCLUSION

Together, these findings identify activation of developmental Notch–WNT signaling as a central mechanism in pediatric iDCM. This signaling axis drives maladaptive remodeling and myocardial stiffening and is functionally targetable *in vivo*. Recognition of this age-specific biology provides a mechanistic framework for developing therapies tailored to pediatric heart failure rather than extrapolated from adult disease.

## LIMITATIONS

Our model of disease represents an early onset model of iDCM as the intent is to investigate disease mechanisms prior to end stage HF and before the rats reach puberty. Although our studies suggest Notch inhibition may be a sound approach to prevent the disease process, the effect of inhibiting β-catenin is yet to be investigated. Furthermore, the molecular mechanism by which chronic sFRP1 switches from WNT antagonist to agonist requires further investigation, potentially involving receptor dynamics or sFRP1 multimerization. Additionally, the mechanisms linking Notch-WNT activation to stiffness (cytoskeletal vs. ECM proteoglycans) require further dissection using cardiomyocyte-specific proteomics and mechanical profiling. A limitation is that our snRNA-seq was generated from one pooled LV sample per group, precluding inference of between-animal variability and increasing risk of pseudo-replication if nuclei are treated as independent replicates. Accordingly, we interpreted the snRNA-seq data as providing a high-resolution map of cell states and pathways within each condition, and we use them to generate mechanistic hypotheses that are supported by bulk RNA-seq and protein-level validation (Notch/β-catenin activation, ECM proteoglycan enrichment).

While global Notch blockade is unlikely to be clinically feasible due to systemic effects, these findings highlight downstream effectors of developmental signaling as potential pediatric-specific therapeutic targets.

## Abbreviations

### A

ACEi: Angiotensin Converting Enzyme Inhibitor
ADAM: A Disintegrin And Metalloproteinase
AFM: Atomic Force Microscopy
AHA: American Heart Association
ANP / ANF: Atrial Natriuretic Peptide / Factor (Protein product of *NPPA*)
ANOVA: Analysis of Variance

### B

β-AR: Beta-Adrenergic Receptor
BNP: Brain Natriuretic Peptide (Protein product of *NPPB*)

### C

CM: Cardiomyocyte
CTNNB1: Catenin Beta 1 (Gene encoding β-catenin)

### D

DAPT: N-[N-(3,5-Difluorophenacetyl)-L-alanyl]-S-phenylglycine t-butyl ester (γ-secretase inhibitor)
DCM: Dilated Cardiomyopathy
DEG: Differentially Expressed Gene
DMSO: Dimethyl Sulfoxide

### E

ECM: Extracellular Matrix
EF: Ejection Fraction
ELISA: Enzyme-Linked Immunosorbent Assay

### F

FB: Fibroblast
FGP: Fetal Gene Program

### G

GAPDH: Glyceraldehyde 3-phosphate dehydrogenase
GDMT: Guideline-Directed Medical Therapy

### H

HF: Heart Failure

### I

iDCM: Idiopathic Dilated Cardiomyopathy
IP: Intraperitoneal
IQR: Interquartile Range
Is: ISO + sFRP1 combined treatment group
ISO: Isoproterenol

### L

LV: Left Ventricle / Left Ventricular
LVAW: Left Ventricular Anterior Wall
LVAW(s): Left Ventricular Anterior Wall in Systole
LVID: Left Ventricular Internal Diameter
LVID(d): Left Ventricular Internal Diameter in Diastole
LVID(s): Left Ventricular Internal Diameter in Systole
LVPW: Left Ventricular Posterior Wall
LVPW(s): Left Ventricular Posterior Wall in Systole
LV vol: Left Ventricular Volume
LV vol(d): Left Ventricular Volume in Diastole
LV vol(s): Left Ventricular Volume in Systole

### M

MHC: Myosin Heavy Chain
MYH6: Myosin Heavy Chain 6 (α-MHC)
MYH7: Myosin Heavy Chain 7 (β-MHC)

### N

NF: Non-Failing (Control)
NICD: Notch Intracellular Domain
NPPA: Natriuretic Peptide A
NPPB: Natriuretic Peptide B
NRVMs: Neonatal Rat Ventricular Myocytes

### O

OCT: Optimal Cutting Temperature compound

### P

PBS: Phosphate Buffered Saline
PDE3i: Phosphodiesterase 3-Inhibitor

### R

RCT: Randomized Controlled Trial
RNA-seq: RNA Sequencing
RT-qPCR: Reverse Transcription Quantitative Polymerase Chain Reaction

### S

SEM: Standard Error of the Mean
sFRP1: Secreted Frizzled Related Protein 1
siRNA: Small Interfering RNA
snRNA-seq: Single-Nuclei RNA Sequencing

### T

TGF: Transforming Growth Factor

### U

UMAP: Uniform Manifold Approximation and Projection

## Acknowledgments

We would like to acknowledge the Heart Transplant Team at Children’s Hospital Colorado, the University of Colorado hemodynamics core, especially Elizabeth Hardy, Juliana Sucharov for aiding in analysis of the single nuclei RNA seq. and the Integrated Physiology Program at the University of Colorado, particularly Dr. Mary Weiser-Evans, the program Director.

## Author contributions

OON performed experiments, data analysis and interpretation, manuscript writing and editing. ER performed experiments, data analysis and manuscript writing. JG performed experiments and data analysis. AKF performed statistical and bioinformatics analyses. CC performed experiments. JCC and MRB were involved in the adult tissue collection and selection. LHL performed experiments. MPAB performed data analysis. BP performed experiments and guided AFM analysis. SDM was involved in experimental design, project supervision, clinical data interpretation, and manuscript writing and editing. BLS was involved in experimental design, project supervision, data interpretation, and manuscript writing and editing. CCS conceptualized the study, was involved in experimental design, data analysis and interpretation, manuscript writing and editing. All authors reviewed and approved the manuscript.

## Data Availability

The datasets supporting the current study including raw, processed and metadata of RNA sequencing will be made publicly available in NCBI GEOdatabase upon acceptance of the manuscript for publication.

## Novelty and Significance

### What is known?

- Pediatric iDCM responds poorly to adult heart failure therapies.
- Developmental pathways are transcriptionally enriched in pediatric iDCM. What new information does this article contribute?
- Notch–WNT co-activation drives pediatric remodeling and stiffness.
- Notch inhibition improves cardiac function *in vivo*.

### Summary

Pediatric iDCM is a biologically distinct disease driven by synergistic activation of developmental Notch and WNT pathways, instead of adult-like fibrotic remodeling. The study uncovers Notch/WNT-mediated stiffness as a pediatric-specific therapeutic target and provides a rational for designing child-focused interventions instead of extrapolating adult guidelines.

