## Supplemental material for "Synergistic Notch–WNT Activation Underlies Pediatric Dilated Cardiomyopathy"

SUPPLEMENTARY MATERIAL

**List of supplementary material**

Detailed Methods

Supplemental tables S1-S6

Supplemental figure and legends S1-S17

References ##58

**DETAILED METHODS**

**Human sample collection**

Heart samples of DCM and NF control patients for both children (n= 7 for NF, n=12 for DCM) and adults (n=4/group) were included in the study for pediatric samples. Patients had a clinical diagnosis of iDCM as documented in the electronic medical record, ejection fraction ≤ 45% and left ventricular end diastolic dimension z score ≥ to 2. NF heart samples were collected from children and adults who had normal cardiac structure, no history of heart disease, were designated as brain dead heart donors at the time of tissue collection, and whose heart could not be placed due to lack of an appropriate recipient (i.e. blood type or size mismatch). During cardiac transplantation, the explanted hearts were immediately cooled in ice cold oxygenated Tyrode’s solution in the operating room. The LV tissue was immediately dissected, flash frozen and stored at -80^0^C until further use. NF subjects and DCM patients were selected based on matched age and sex. Detailed patient demographics are listed in Supplemental Table 1. Clinical characteristics and medications reflect time of tissue collection. Human tissue collection was approved by the University of Colorado Anschutz Medical Campus-Colorado Multiple Institutional Review Board (COMIRB) and complies with the Declaration of Helsinki. Written informed consent was obtained from each patient, parent or guardian. Donor tissue collection was consented by family members.

**Cell culture and transfection**

NRVMs were prepared from 1 to 3-day old rats as extensively described by our group^9^ and were treated for 72 hours prior to harvesting. Treatments were as follows: 1µg/ml sFRP1 (Recombinant Human sFRP-1 Protein, CF, R&D systems), 100nM ISO (Sigma Aldrich), and 20µM DAPT (Sigma Aldrich), a Notch receptor antagonist. NRVMs were transfected with 20nM siRNAs for *CTNNB1* (Thermofisher, s438) or scrambled control using RNAiMAX (Invitrogen) as we described^9^.

**RNA extraction and real-time quantitative PCR.**

NRVMs were homogenized in Qiazol (QIAGEN), and RNA was extracted and precipitated using the chloroform/isopropanol method optimized by our laboratory^9,44^. cDNA was synthesized using the Applied Biosystems High-Capacity cDNA Synthesis Kit (Thermo Fisher Scientific) according to manufacturer’s instructions. Gene expression was measured by real-time quantitative PCR (RT-qPCR) as previously described^9^, with Power Sybr Green PCR Master Mix (Life Technologies). Expression levels of all transcripts were normalized to 18S rRNA, and no difference in the expression of 18S between groups was appreciated (data not shown). RT-qPCR primer sequences^45^ are listed in Supplemental Table 5.

**Protein extraction and nuclear/cytoplasmic fractionation**

Total protein was extracted from cells using RIPA40 (Pierce RIPA Buffer, FisherScientific, PI89900). Nuclear and cytoplasmic fractions were purified using NE-PER (FisherScientific, 78835) as described by our group^9,16,46,47^. Immunoblots were performed essentially as previously described with minor modification^45,46^. Briefly, transfer was performed using the Trans-Blot turbo transfer system (Biorad 1704150), blocked with 5%BSA in PBS-T for 1 hour and incubated with primary antibody overnight at 4^0^C and with secondary antibody for 1 hour at room temperature. Chemiluminescence was captured using High-sensitivity Chemiluminescence with image lab software (Biorad Chemi Doc Imaging system 12003153). Antibodies used for Western blot analysis are listed in Supplemental Table 6.

**Neonatal rat injections**

All animal studies were in compliance with the *Guide for the Care and Use of Laboratory Animals* (National Academies Press, 2011) and approved by the IACUC of the University of Colorado Anschutz Medical Campus. Animal protocols are in accordance with Public Health Service (PHS) Animal Welfare Assurance (A3269-01). Pregnant Sprague Dawley female rats were purchased from Charles Rivers laboratories. Animals were housed in the animal facility on Campus and monitored daily. Day 5 neonatal rats were injected intraperitoneally (IP) with 0.1mg/kg/day ISO every other day (dose previously optimized by our group and others and reflects circulating catecholamine levels in children)^48,49^ for 5 treatments and every day for the subsequent 7 days. sFRP1 (50µg/kg/day) (dose previously optimized by our group)^6,9,50^ was co-administered with ISO (or vehicle (PBS, 0.5mM ascorbic acid). DAPT (500µg/kg/day) (dose optimized by another group)^51^ or vehicle was administered con-currently with ISO+sFRP1 and was dissolved in 40%PEG300, 5%Tween 80, 50%ddH2O, and 5%DMSO. Injections were performed in male and female animals. The sFRP1 (Recombinant Human sFRP-1 Protein, CF, R&D systems) solution was freshly prepared on the days of treatment by dissolving it in phosphate-buffered saline (PBS) at room temperature. ISO (Isoproterenol hydrochloride, Sigma Aldrich St. Louis, MO) was freshly prepared and dissolved in 0.5mM ascorbic acid.

**Echocardiographic assessment**

Transthoracic echocardiograms (EchoPAC SW v201) were performed to evaluate cardiac function. Views were taken in planes which approximated the parasternal short axis and long axis views in humans. LV internal diameters and wall thicknesses were measured (average of at least 3 cardiac cycles) at end-systole and end-diastole^47^. Morphometrics were collected at the time of sacrifice and hearts were frozen at -80ºC. Measurements were performed by the University of Colorado hemodynamics core.

**Isolation of myocardium**

At the end of the study period, rats were euthanized, myocardial tissue was isolated, immediately weighed, frozen in liquid nitrogen and stored at -80°C for further molecular analysis. Sections of myocardium from left ventricular (LV) tissue were evaluated by Hematoxylin and Eosin (H&E) and/or trichrome staining to evaluate for fibrosis and hypertrophy as described^6,16,52^. All slides were evaluated and analyzed in a blind fashion. The percentage area of fibrosis for trichrome and H&E staining was analyzed with ImageJ. To analyze cardiomyocyte hypertrophy, cell area and number were quantified as previously described by our group^6^.

**AFM assessment of stiffness**

AFM assessment of cardiac tissue was done as extensively described by our group^53,54^. Briefly, flash frozen myocardial tissue was embedded with optimal cutting temperature (OCT) compound. Tissues were allowed to equilibrate at cryostat temperature of -20°C and cryosectioned at 5 microns. The morphological details of isolated myocardial tissue on slides were observed with an optical light microscope. We determined tissue stiffness using a NanoWizard® 4a (JPK Instruments, Carpinteria, CA, USA). Tissue was scanned in quantitative imaging (QI) mode with a qp-BioAC-1 (NanoandMore, Watsonville, CA, USA) cantilever. The cantilever was calibrated prior to each experiment using the thermal oscillation method with a force constant in the range of 0.15 to 0.55 N/m. A 5,625 µm^2^ area was scanned using a set point of 5 nN and a Z-length of 2 µm. At least four random scans were performed per tissue sample with several tissue orientations scanned across the sample. These scans (each individual scan composed of over 60,000 force curves) were considered for the mechanical average of each sample. Data was analyzed with JPK software. The Hertz model was used to determine the mechanical properties of the myocardium using the JPK software at the Bioscience department of the University of Colorado.

**Bulk RNA-sequencing**

Total RNA was extracted from rats injected with ISO, sFRP1, ISO+sFRP1 and vehicle-treated controls (n=6/group), quantified using Qubit (Biotium) fluorometric quantitation, and assessed for quality using the Agilent Bioanalyzer Nano RNA Chip (Agilent)^55^. Samples with an RNA integrity number greater than 9 were considered to be high quality and suitable for RNA-Seq.

RNA sequencing reads were obtained using the Illumina HiSeq analysis pipeline. Read quality was confirmed using FastQC ([**http://www.bioinformatics.**](http://www.bioinformatics.bbsrc.ac.uk/projects/fastqc#_blank)). All adapter sequences were trimmed from raw sequencing files, and the reads were mapped to the norvegicus genome (Rnor_6.0) using STAR^55^. Counts estimates were calculated using FeatureCounts^56^. Counts were normalized using edgeR. Wilcoxon rank-sum test was used to compare groups on normalized data, and a heatmap was generated for significant results (p-value<0.05 and fold change>1.75)

**Single nuclei RNA-Sequencing and nuclei isolation**

Nuclei were isolated from LV tissue of three rats per condition pooled to generate one library per group. Each condition was therefore represented by one aggregated snRNA-seq dataset (pseudobulk at the sample level), and analyses were focused on identifying cell populations and pathway-level patterns rather than performing differential expression testing across independent biological replicates. The following groups were analyzed: ISO, sFRP1, ISO+sFRP1 or vehicle-treated controls using the Chromium Nuclei Isolation Kit (10x Genomics, PN-1000494) and as described extensively by the Lavine lab^18^**.** Reverse transcription, barcoding, complementary DNA amplification and purification for library preparation were performed by the Genomics Core at CU Anschutz using the 10X Genomics Chromium technology to capture and profile single nuclei transcriptome 3’ gene expression. Generated libraries were sequenced on the Illumina NovaSeq 6000 instrument at the Genomics and Microarray Core of University of Colorado Anschutz Medical Center. Upon sequencing, Fastq sequencing files were processed through Cell Ranger (v7.1.0). Data processing downstream was performed in the R package Seurat (v4)^57^. Despite most cells falling below these thresholds, both mitochondrial reads/cell and ribosomal reads/cell had a threshold set to <10%, read counts/cell>5000, and read counts/cell<40000 were applied. Following filtering, normalization and clustering (cca_clusters) followed the standard Seurat pipeline. Azimuth (https://app.azimuth.hubmapconsortium.org/app/human-heart) was used to determine cell identity along with known cell markers^58^ (see supp. Table 3.1 for markers used). Dimensionality reduction was done using Principal Component Analysis (PCA), clusters were annotated into major cell populations, and the ventricular cardiomyocyte cluster was sub-clustered to eight different sub-clusters (supp. Table 3.3) and data visualized through UMAP projection. Differentially expressed genes for each cluster and sub-cluster were calculated with the FindAllMarkers function the Wilcoxon Rank Sum Test with a minimum fraction (min.pct) of 0.10 and a logFC threshold of 0.15 in R package Seurat (v4) (supp. Table 3.2).

**Intercellular Communication Analysis**

Mapping of the cardiac interactome and cell-to-cell communication was conducted using the CellChat framework to compare signaling patterns between conditions. CellChat was applied to each condition as a single dataset to infer and compare intercellular communication networks, recognizing that inferences are based on one sample per condition and thus should be interpreted as hypothesis-generating rather than definitive across animals. We quantified the global communication landscape by calculating the differential number of interactions and differential interaction strength. Additionally, to identify pathological signaling shifts, we focused on the significant networks such as NOTCH signaling.

The analysis included:

1. Signaling Pattern Recognition: We identified incoming and outgoing signaling patterns to determine how specific cell types, such as smooth muscle cells and fibroblast, coordinate their communication in the diseased state.
2. Pathway Specificity: Dot plots and scatter plots were utilized to categorize pathways as Shared, Control-specific, or ISO+sFRP1-specific.

**Statistical Comparison of Signaling Networks**

Comparisons of interaction frequency were performed across all cell types to identify the total number of interactions per condition. Network centrality and hierarchy plots were generated for the NOTCH pathway to visualize changes in network density and the directionality of signals between celltypes. Relative signaling strengths were normalized and compared using heatmap visualization of overall signaling patterns

**Pathway Analysis**

Volcano plot and hierarchical clustering/heatmap generation were performed using R (The R Foundation, R Core Team (2023). R: A Language and Environment for Statistical Computing. R Foundation for Statistical Computing, Vienna, Austria.https://www.R-project.org/). Gene set enrichment analysis, Reactome and Enrichr pathway analysis platforms were used to investigate biological and clinically relevant pathways associated with significantly dysregulated genes.

**Statistics**

Statistical analyses for all omics data are described in each omics section. All other analyses (RT-qPCR, western blot and AFM) were performed using GraphPad Prism 8 (GraphPad Software), and significance threshold is set a priori at *p*<0.05. AFM data were checked for normality using Shapiro-Wilk. If not normally distributed, the Mann-Whitney *U* test was used. Comparisons between two groups were made using unpaired two-tailed Student’s *t*-test. For comparisons involving more than two groups, one-way ANOVA with Tukey’s post hoc test was used as appropriate Quantitative results are presented as mean ± SEM. Each appropriate statistical test is reported in the figure legends.

**SUPPLEMENTAL TABLE LEGENDS**

**Supp. Table 1. Detailed patient characteristics.**

| **ID** | **Group** | **Sex** | **Age yrs** | **EF %** | **Medical device** | **PDE3i** | **PDE5i** | **Non PDEi ionotrope** | **Digoxin** | **ACEi/ ARBs** | **Beta blocker** | **Diuretic** | **Western blot** | **AFM** |
| --- | --- | --- | --- | --- | --- | --- | --- | --- | --- | --- | --- | --- | --- | --- |
| 1 | NF | M | 1.4 | NA | No |  |  |  |  |  |  |  | X | X |
| 2 | NF | M | 9.5 | NA | No |  |  | X |  |  | X |  | X |  |
| 3 | NF | M | 12 | NA | No |  |  | X |  |  |  | X | X |  |
| 4 | NF | M | 7 | NA | No |  |  |  |  |  |  |  | X |  |
| 5 | NF | F | 3.2 | NA | No |  |  | X |  |  |  |  | X |  |
| 6 | NF | F | 8 | NA | No |  |  |  |  |  |  |  | X | X |
| 7 | NF | M | 7 | NA | No |  |  | X |  |  |  |  | X | X |
| NF | n=7 | 28.6% F | 6.9 | 0.0% | 0.0% | 0.0% | 0.0% | 57.1% | 0.0% | 0.0% | 14.3% | 14.3% | 100% | 42.9% |
| 8 | iDCM | F | 3.6 | 28.00 | No | X |  |  |  | X |  | X | X |  |
| 9 | iDCM | F | 0.8 | 35.00 | No |  |  |  |  | X |  | X | X |  |
| 10 | iDCM | F | 4.1 | 12.00 | ECMO | X |  | X |  | X |  |  | X |  |
| 11 | iDCM | M | 7 | 20.00 | VAD (Berlin) | X | X | X |  | X |  | X | X |  |
| 12 | iDCM | F | 6.6 | 11.00 | No | X |  |  |  | X |  | X | X |  |
| 13 | iDCM | M | 3.12 | 13.00 | No |  |  | X |  | X |  | X | X |  |
| 14 | iDCM | M | 3.04 | 14.00 | Pacemaker, balloon valvularplasty | X |  |  | X | X |  | X | X |  |
| 15 | iDCM | F | 4.4 | 13.00 | Pacemaker |  |  |  |  | X |  | X | X |  |
| 16 | iDCM | F | 0.77 | 8.00 | No | X |  | X | X | X |  | X | X | X |
| 17 | iDCM | M | 2.82 | 15.00 | No | X |  |  |  | X |  | X | X | X |
| 18 | iDCM | M | 1 | 17.00 | No | X |  |  |  | X |  | X | X | X |
| 19 | iDCM | F | 10.48 | 22.00 | No | X |  | X | X | X | X | X |  | X |
| iDCM | n=12 | 58.3%F | 3.9 | 17.3% | 33.3% | 75% | 8.3% | 41.7% | 25% | 100% | 8.3% | 91.7% | 91.7% | 33.3% |
| 20 | NF | M | 52 | NA | No |  |  |  |  |  | X | X | X |  |
| 21 | NF | F | 56 | NA | No |  |  |  |  |  |  |  | X |  |
| 22 | NF | F | 61 | NA | No |  |  |  |  |  |  | X | X |  |
| 23 | NF | M | 46 | NA | No |  |  |  |  |  |  |  | X |  |
| NF | n=4 | 50% F | 55.2 | 0.0% | 0.0% | 0.0% | 0.0% | 0.0% | 0.0% | 0.0% | 25% | 50% | 100% |  |
| 24 | iDCM | M | 52.8 | 10 | No |  |  | X |  | X |  |  | X |  |
| 25 | iDCM | F | 57.6 | 13.75 | BiVICD |  |  | X | X |  |  | X | X |  |
| 26 | iDCM | F | 59 | NA | No |  |  |  |  |  |  | X | X |  |
| 27 | iDCM | M | 39.8 | 11 | IABP, AICD | X |  | X | X | X |  | X | X |  |
| iDCM | n=4 | 50% F | 54 | 11.58 | 50% | 25% | 0.0% | 75% | 50% | 50% | 0.0% | 75% | 100% |  |
| Table S1: Patient characteristics. This includes the detailed deidentified patient characteristics including sex, age at tissue collection, EF, medical device use, medication use, and experiments done for each sample. Median age for pediatric iDCM patients was 3.9 years (interquartile range-IQR=6.2), pediatric non-failing control (NF) subjects had a median age of 6.9 years (IQR=9.5). Median age for adult iDCM was 54 years (IQR=15). Adult NF patients had a median age of 55.2 years (IQR=19.2),. PDE3i, PDE5i, ACEi and diuretics are more commonly used in DCM patients. #Inotropes include: dobutamine, dopamine, epinephrine, norepinephrine. *Last EF prior to mechanical assist device placement.  NF = Non-failing, ID = Identification, DCM = Dilated cardiomyopathy M = Male, F = Female, EF = Ejection fraction, NA=Not applicable, BiVICD= Biventricular Implantable Cardioverter-Defibrillator, IABP= Intra-aortic Balloon Pump, AICD= Automatic Implantable Cardioverter-Defibrillator, ACEi = Angiotensin-converting enzyme inhibitor, PDE3i = Phosphodiesterase inhibitor 3, Other inotropes include: dobutamine, dopamine, epinephrine, norepinephrine, and vasopression. | | | | | | | | | | | | | | |

**Supp. Table 2. Echocardiogram findings of rats treated with ISO, sFRP1 and ISO+sFRP1 compared to Vehicle treated controls.**

| Tukey's multiple comparisons test | EF – mean difference/ p-value | LV vol(s) – mean difference/ p-value | LVID(s) – mean difference/ p-value | LVAW(s)- mean difference/ p-value | LVPW(s)- mean difference/ p-value | LV vol(d) – mean difference/ p-value | LVID(d) – mean difference/ p-value |
| --- | --- | --- | --- | --- | --- | --- | --- |
| Control vs. ISO | 4.590/  0.1388 | -4.920/  0.7825 | -0.2312/  0.5797 | 0.05847/  0.8184 | 0.05379/  0.8076 | -1.779/  0.9974 | -0.1745/  0.7937 |
| Control vs. sFRP1 | -0.1634/  0.9998 | 2.222/  0.9738 | 0.1422/  0.8597 | 0.02892/  0.9932 | 0.01802/  0.9991 | 13.39/  0.4687 | 0.2246/  0.6385 |
| Control vs. ISO+ sFRP1 | 10.58/  <0.0001 | -20.04/  0.0002 | -0.7456/  <0.0001 | 0.01962/  0.9978 | 0.06735/  0.3722 | -21.25/  0.0108 | -0.5327/  0.0030 |
| Echocardiographic assessment of left ventricular function in neonatal rats following treatment with Isoproterenol (ISO), secreted frizzled protein 1 (sFRP1), or combined ISO+sFRP1 compared with vehicle controls. Parameters include ejection fraction (EF), left ventricular volume (LV vol), left ventricular internal diameter (LVID), left ventricular anterior (LVAW) and posterior wall (LVPW) both in systole (s) and diastole(d). Data are expressed as mean ± SEM/p-value. Statistical significance was determined using one-way ANOVA with post hoc analysis as described in the Methods | | | | | | | |

**Supp. Table 3. Differential gene expression from Bulk RNA sequencing in left ventricular tissue from Vehicle (control) and ISO+sFRP1-treated rats.**

Table includes gene names, log_2_ fold change, p-values_ttest for each gene. Counts estimates were calculated using FeatureCounts. Counts were normalized using edgeR. The Wilcoxon rank-sum test was used to compare groups on normalized data. n = 6 per group. (Attached as large data set).

**Supp. Table 4. Single nuclei RNA sequencing in left ventricular tissue from Vehicle (control), ISO-, sFRP1- and ISO+sFRP1- treated rats.**

List of canonical marker genes used to assign cellular identities to nuclei clusters obtained from snRNA-seq of LV tissue from Vehicle (control), ISO-, sFRP1- and ISO+sFRP1- treated rats. Marker genes (Supp. Table 4.1) were selected based on published literature and using the Azimuth database. The table includes gene names, average log_2_ fold change and p-values for each cluster. DEGs (Supp. Table 4.2) identified in the ventricular cardiomyocyte cluster of ISO+sFRP1 treated rats compared to vehicle control. The table includes gene names, average log_2_ fold change, and p-values for each gene. Summary of all cell clusters identified (Table 4.3). Each cluster is described as number of cells, percentage of total cells. n = 1 per group (3 rats combined per group). (Attached as large data set).

**Supp. Table 5. Primer sequences used for Real time-quantitative polymerase chain reaction.**

| **Rat Primers** | **Forward sequence** | **Reverse sequence** |
| --- | --- | --- |
| *Notch 1* | 5’ ATACGCCTGTGGCAGAATAAG 3’ | 5’ CGGGACAGACTTGTTCCTTTAG 3’ |
| *Notch 2* | 5’ TGGAGGTCTCAGTGGCTATAA 3’ | 5’ TTCTGGCACGGGTTAGAAAG 3’ |
| *Notch 3* | 5’ GCTTGATTTCCCATACCCACTA 3’ | 5’ CGAAGATGACCAGCAGAAAGA 3’ |
| *Notch 4* | 5’ GGTGGAAGGAATCAGAACTAGG 3’ | 5’ GGGAAGACAAGTGGGCATAA 3’ |
| *HEY 1* | 5’ CTGCAGGAGGGAAAGGTTATT 3’ | 5’ CTCAGATAACGGGCAACTTCA 3’ |
| *HEY2* | 5’ AGATCTCCACAGCAGCAATAAA 3’ | 5’ GTTGCCAAGCTGCCTTAAAC 3’ |
| *HES1* | 5’ TCAAAGCCTATCATGGAGAAGAG 3’ | 5’ GAATGCCGGGAGCTATCTTT 3’ |
| *HEYL* | 5’ ACCACCATCCTCCAGAACTA 3’ | 5’ GGGCTGAATCCCAAGAGAA 3’ |
| *sFRP1* | 5’ GGTCAAGCCCAGAAAGTGATA 3’ | 5’ AAAGACTGTGGGCAGAGAAG 3’ |
| *Col1α1* | 5’ CGGACTATTGAAGGAGCCTAAC 3’ | 5’TACACAAGGAACAGAACAGTCTC3’ |
| *COL3α* | 5’ ATGTGGGACCTGGTTTCTTC 3’ | 5’ CAGTCTAGTGGCTCATCATCAC3’ |
| *TGFβ1* | 5’CCAGATCCTGTCCAAACTAAGG 3’ | 5’ TTGTTGCGGTCCACCATTA 3’ |
| *galectin-3* | 5’ CTGACAGTGCCCTACGATATG 3’ | 5’ GTTATTGTCCTGCTTCGTGTTG 3’ |
| *Corin* | 5’ GTTATTGTCCTGCTTCGTGTTG3’ | 5’ GGAACTTCTCCGTGTCCATATT 3’ |
| *matrix metalloproteinase-9* | 5’ GCTGCTCCAACTGCTGTATAA 3’ | 5’ TGGTGTCCTCCGATGTAAGA 3’ |
| *matrix metalloproteinase-2* | 5’ CACCAAGAACTTCCGACTATCC 3’ | 5’ TCCAGTACCAGTGTCAGTATCA 3’ |
| *TIMP 1* | 5’ TGGCATCCTCTTGTTGCTATC3’ | 5’ CCACAGCGTCGAATCCTTT 3’ |
| *TIMP 2* | 5’ ACCTGACAAGGACATCGAATTTA 3’ | 5’ CCATCTGGTACCTGTGGTTTAG 3’ |
| *Timp3* | 5’ GTGTCAAGTTTCCTCTGGGTAG3’ | 5’ GTAGATCAGGAGCGCCATATTAG 3’ |
| *TIMP 4* | 5’ TATGGAAGGTGGGCTGACTA 3’ | 5’ CTGCCAGGACAAGTATCAGAAG 3’ |
| *CTNNB1* | 5’ CTCAGATGGTGTCTGCCATAG 3’ | 5’ TGGTGGGAAAGGTTGTGT 3’ |
| Table S5. List of forward and reverse primer sequences used for quantitative RT-qPCR validation of gene expression in rat NRVMs and Rat LV tissue. Primer sequences were designed using PRIMER-BLAST or obtained from previously published data from our group. Products were verified by melt curve analysis. | | |

**Supp. Table 6. Primary antibodies used for immunoblotting.**

| **Antibody** | **Vendor** | **Catalogue number** |
| --- | --- | --- |
| β-Catenin Antibody | Cell signaling | 9562 |
| Notch1 (D1E11) XP® Rabbit mAb | Cell signaling | 3608 |
| GAPDH-HRP | Cell signaling | 3683 |
| Histone 3 antibody | Cell signaling | 9715 |
| Table S5. List of primary antibodies used for immunoblotting analysis of protein expression. The table includes target protein (antibody used), Vendor and catalogue number. | | |

**SUPPLEMENTAL FIGURES AND LEGENDS**

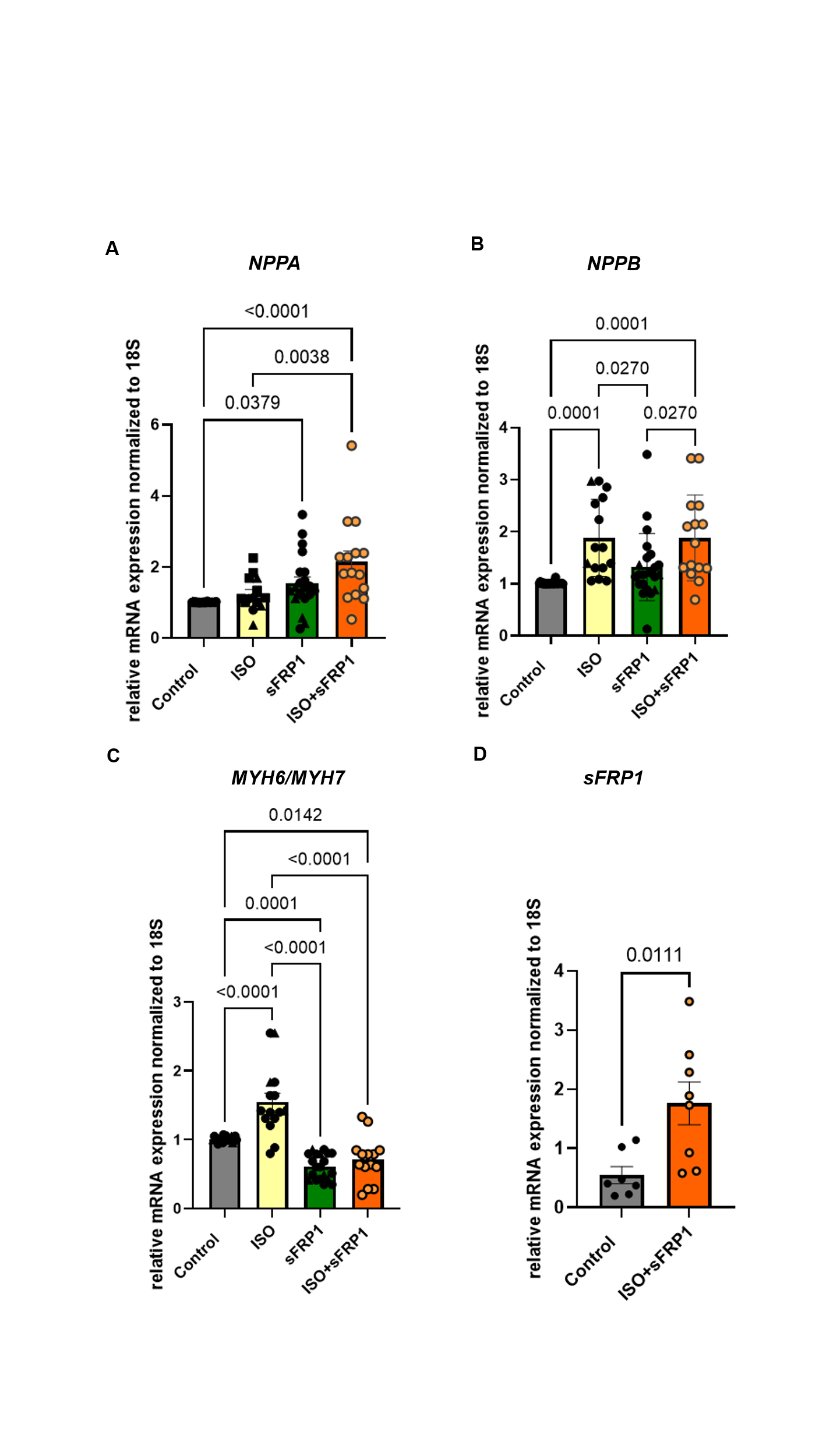

**Supp. Fig. 1.** ISO+sFRP1 promotes pathological remodeling in NRVMs.

(A) RT-qPCR of *NPPA* expression in NRVMs treated with vehicle (Control), ISO, sFRP1 or ISO+sFRP1. Gene expression was normalized to 18S, and data are presented as a relative fold change to Control. NRVMs were treated with 100nM Isoproterenol (ISO) +/- 1 μg/mL human recombinant sFRP1 for 72 hours. n=8 independent NRVM preps. All groups are log_2_ transformed. Fitting a mixed model, Tukey’s multiple comparisons test was used for all data sets.
(B) RT-qPCR of *NPPB* expression in NRVMs treated with vehicle (Control), ISO, sFRP1 or ISO+sFRP1. Gene expression was normalized to 18S, and data are presented as a relative fold change to Control. NRVMs were treated with 100nM Isoproterenol (ISO) +/- 1 μg/mL human recombinant sFRP1 for 72 hours. n=8 independent NRVM preps. All groups are log_2_ transformed. Fitting a mixed model, Tukey’s multiple comparisons test was used for all data sets.
(C) RT-qPCR of *MYH6/7* expression in NRVMs treated with vehicle (Control), ISO, sFRP1 or ISO+sFRP1. Gene expression was normalized to 18S, and data are presented as a relative fold change to Control. NRVMs were treated with 100nM Isoproterenol (ISO) +/- 1 μg/mL human recombinant sFRP1 for 72 hours. n=8 independent NRVM preps. All groups are log_2_ transformed. Fitting a mixed model, Tukey’s multiple comparisons test was used for all data sets.
(D) RT-qPCR of sFRP1 gene expression in NRVMs treated with ISO+sFRP1 compared to vehicle-treated controls. Gene expression was normalized to 18S, and data are presented as a relative fold change to control. n=6 independent NRVM preps. Unpaired 2-tailed t test was used in analysis. All groups are log_2_ transformed and error bar denotes mean± SEM.

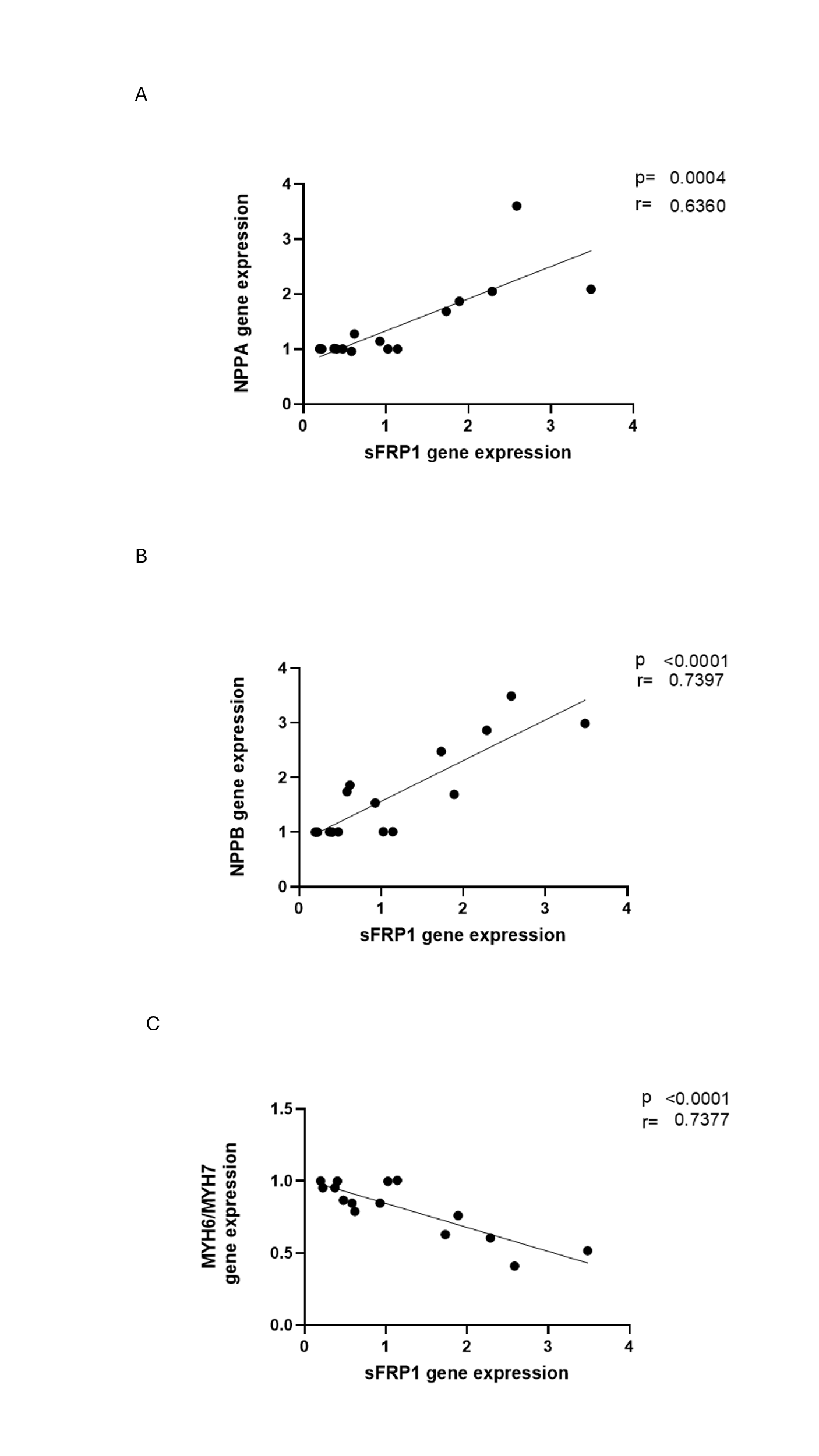

**Supp. Fig. 2. Increase in sFRP1 gene expression in ISO+sFRP1-treated NRVMs correlated with the activation of the FGP**

Simple linear regression of sFRP1 gene expression and expression of (A) *NPPA,* (B) *NPPB* and (C) *MYH6/MYH7*. NRVMs were treated with 100nM Isoproterenol (ISO) +/- 1 μg/mL human recombinant sFRP1 (sFRP1) for 72 hours. n*=*6 independent NRVM preps/groups. *p* values are notated in the figure. Simple linear regression was performed using graph pad prism.

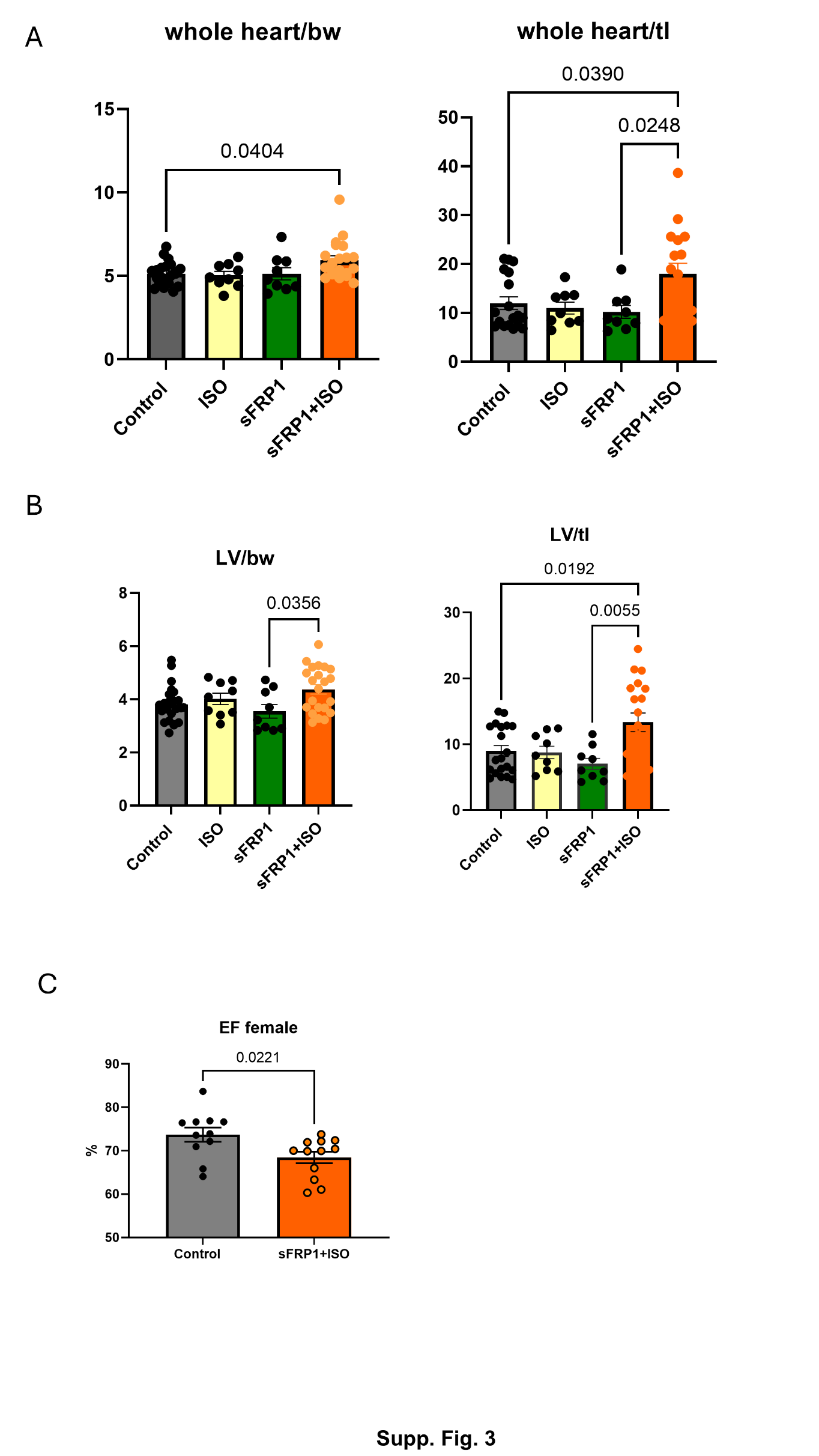

**Supp. Fig. 3. Morphometric data of rats treated with Vehicle control, ISO, sFRPI and ISO+sFRP1**

(A)Morphometric data of whole heart to body weight (left) and whole heart to tibia length (right). Fitting a mixed model, Tukey’s multiple comparisons test was used for all data sets. Control *n=*21, ISO *n* =9, sFRP1 *n* =9 ISO+sFRP1 n=21.

(B) LV weight to body weight (left) and LV weight to tibia length ratios (right) in ISO+sFRP1 rats compared to vehicle-treated controls. Fitting a mixed model, Tukey’s multiple comparisons test was used for all data sets. n=21/group

(C) EF of female neonatal rats treated with Vehicle Control and ISO+sFRP1.

EF, ejection fraction. Control *n=*11, ISO+sFRP1 n=12. Only significant *p*-values are notated in the Figure. Unpaired 2-tailed t test was used in analysis

ISO, Isoproterenol; sFRP1, secreted frizzled protein 1; bw, body weight; tl, tibia length

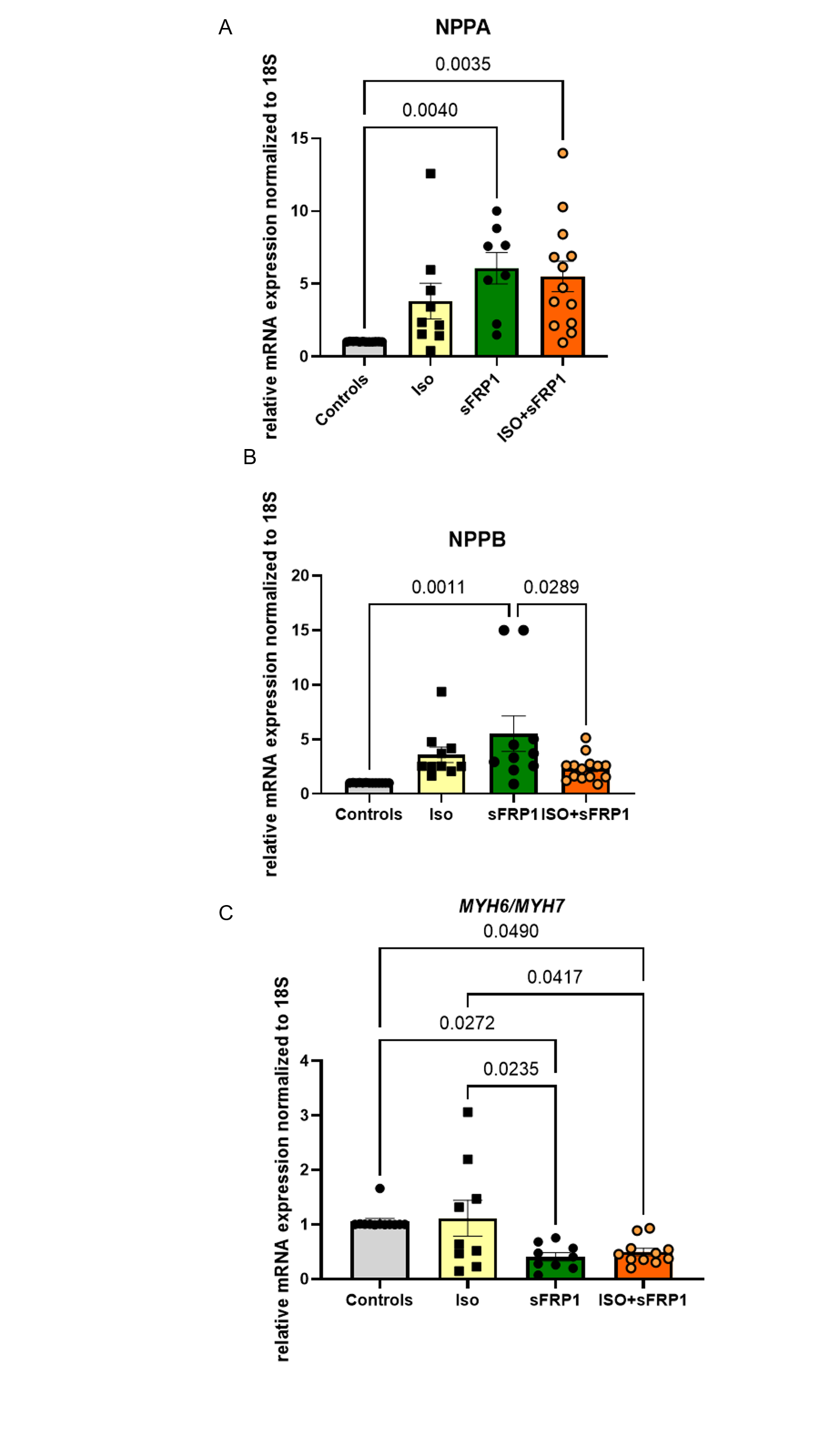

**Supp. Fig 4. Fetal gene expression in rats treated with ISO+sFRP1**

(A-C) RT-qPCR of FGP expression in rats treated with vehicle (Control), ISO, sFRP1 or ISO+sFRP1. Expression of (A) *NPPA*, (B) *NPPB*, and (C) *MYH6* to *MYH7* ratios. Gene expression was normalized to 18S, and data are presented as a relative fold change to Controls. Control *n* = 12, ISO *n* = 9, sFRP1 *n* *= 8*, ISO+sFRP1 *n* = 13. All groups are log_2_ transformed and error bar denotes mean± SEM. Only significant *p* values are notated in the figure. Fitting a mixed model, Tukey’s multiple comparisons test was used for all data sets. ISO, Isoproterenol; sFRP1, secreted frizzled protein 1.

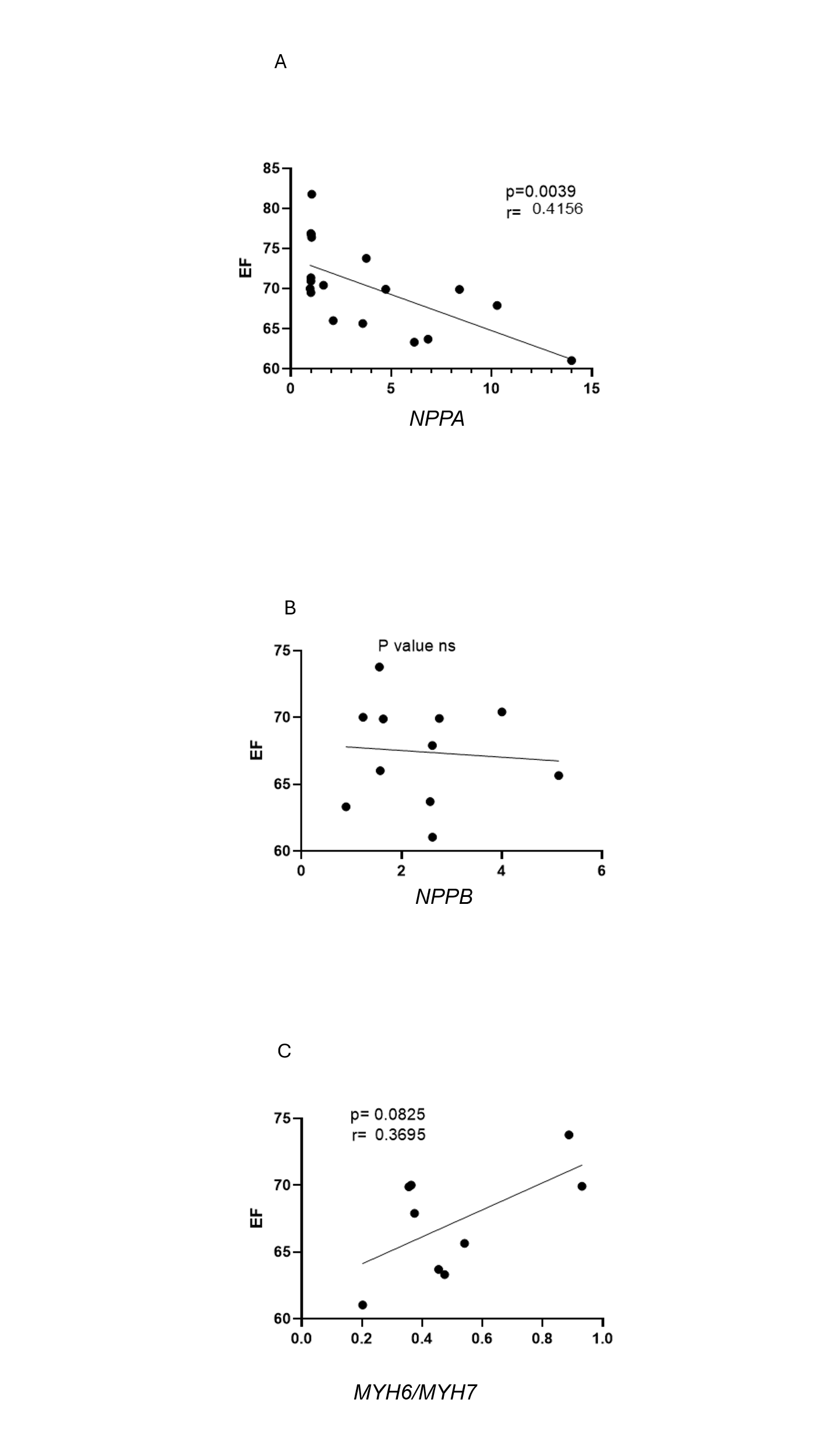

**Supp. Fig. 5. Reactivation of FGP in ISO+sFRP1-treated rats correlates with cardiac dysfunction.**

Simple linear regression of EF and expression of (A) *NPPA,* (B) *NPPB* and (C) *MYH6/MYH7*. *NPPA* *n=15, NPPA, n=11 and MYH6/MYH7* n*=9*. *p* values are notated in the figure.

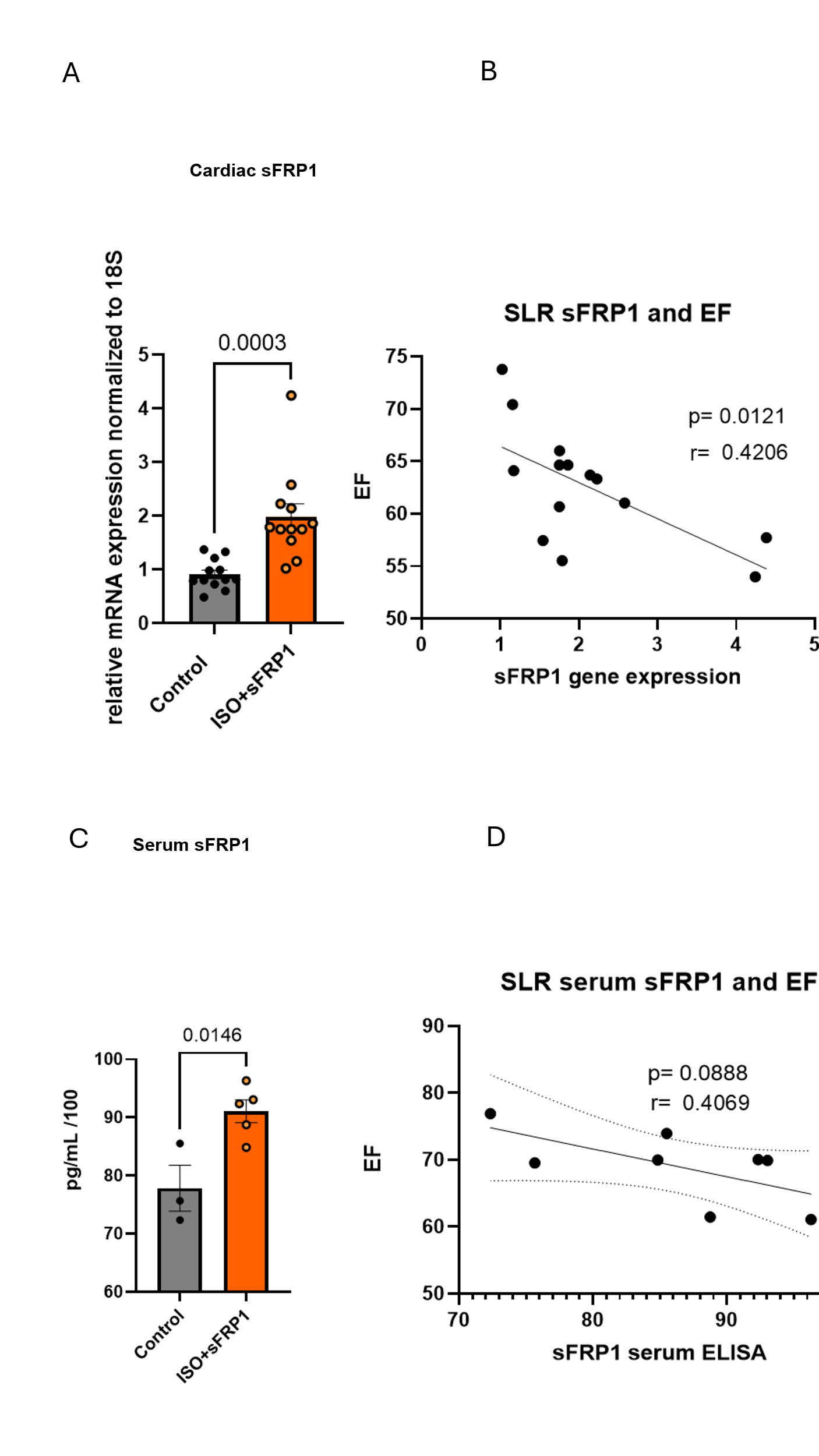

**Supp. figure 6. Higher sFRP1 levels correlate with cardiac dysfunction.**

(A) Levels of sFRP1 mRNA in left ventricle (LV) tissue of ISO+sFRP1-treated rats compared to controls (Left). Gene expression was measured by RT-qPCR and normalized to 18S. n=12/group. Data are presented as a relative fold change to NF controls; all data are log_2_ transformed.

(B) Simple linear regression of sFRP1 gene expression and changes in EF (Right).

(C) Levels of sFRP1 in serum from ISO+sFRP1 rats compared to controls. Serum levels were measured by ELISA. Control n=3, ISO+sFRP1 *n* = 5. ELISA, Enzyme-linked immunosorbent assay.

(D) Simple linear regression of serum sFRP1 levels and changes in EF. Unpaired 2-tailed *t*-test was used in all analyses. Only Significant *p*-values (p<0.05) are notated in the Figure. Error bar denotes mean ± SEM.

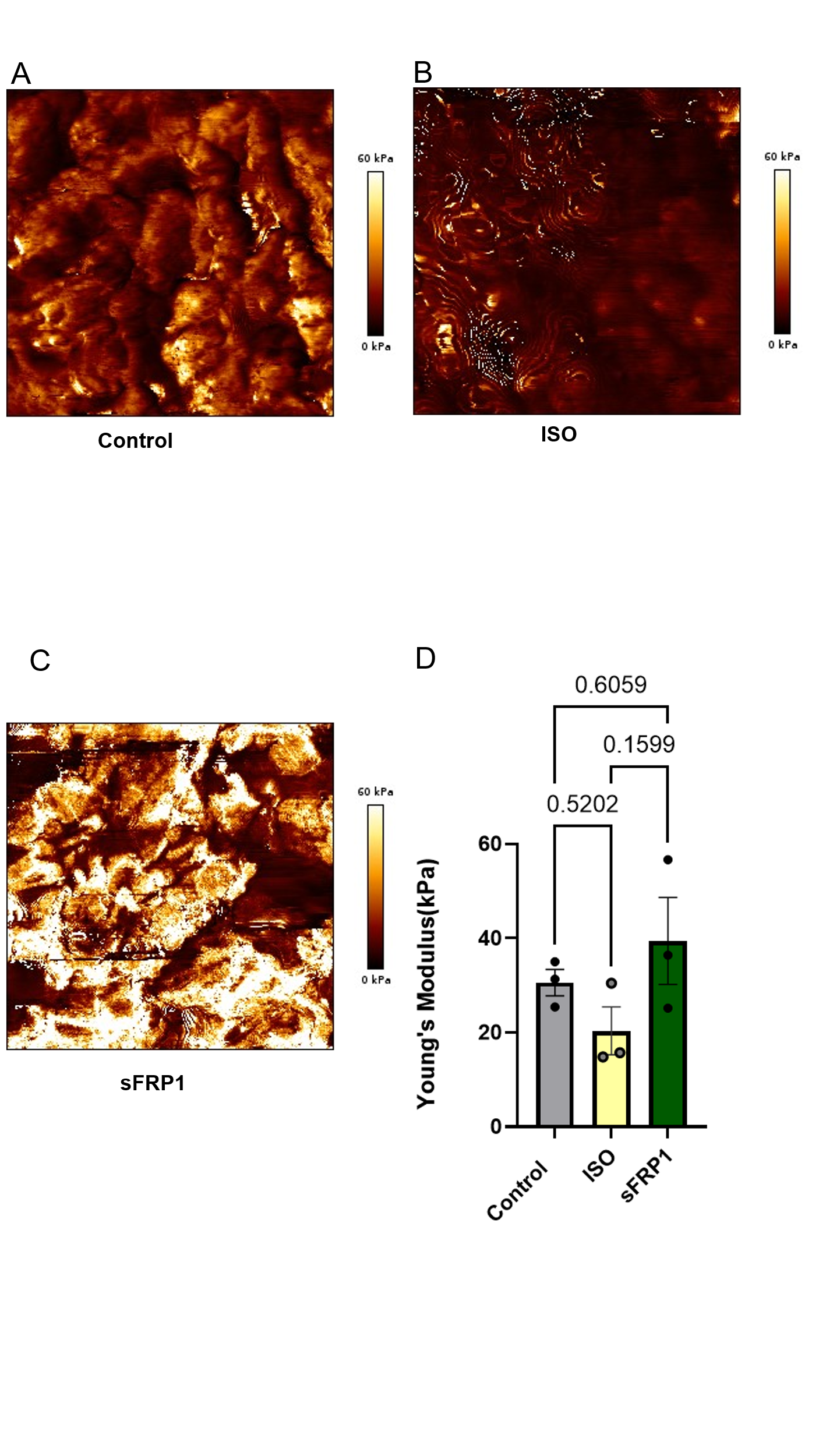

**Supp. Figure 7. Absence of myocardial stiffness in ISO only and sFRP1 only treated rats.**

Elasticity/stiffness (Young’s modulus) in rats treated with ISO (B) or sFRP1 (C) compared to controls (A). Quantification in (D) n=3 per group. Only Significant *p*-values (p<0.05) are notated in the Figure. Tukey’s multiple comparison was used in analyses. Error bar denotes mean ± SEM. ISO, Isoproterenol; sFRP1, secreted frizzled protein 1.

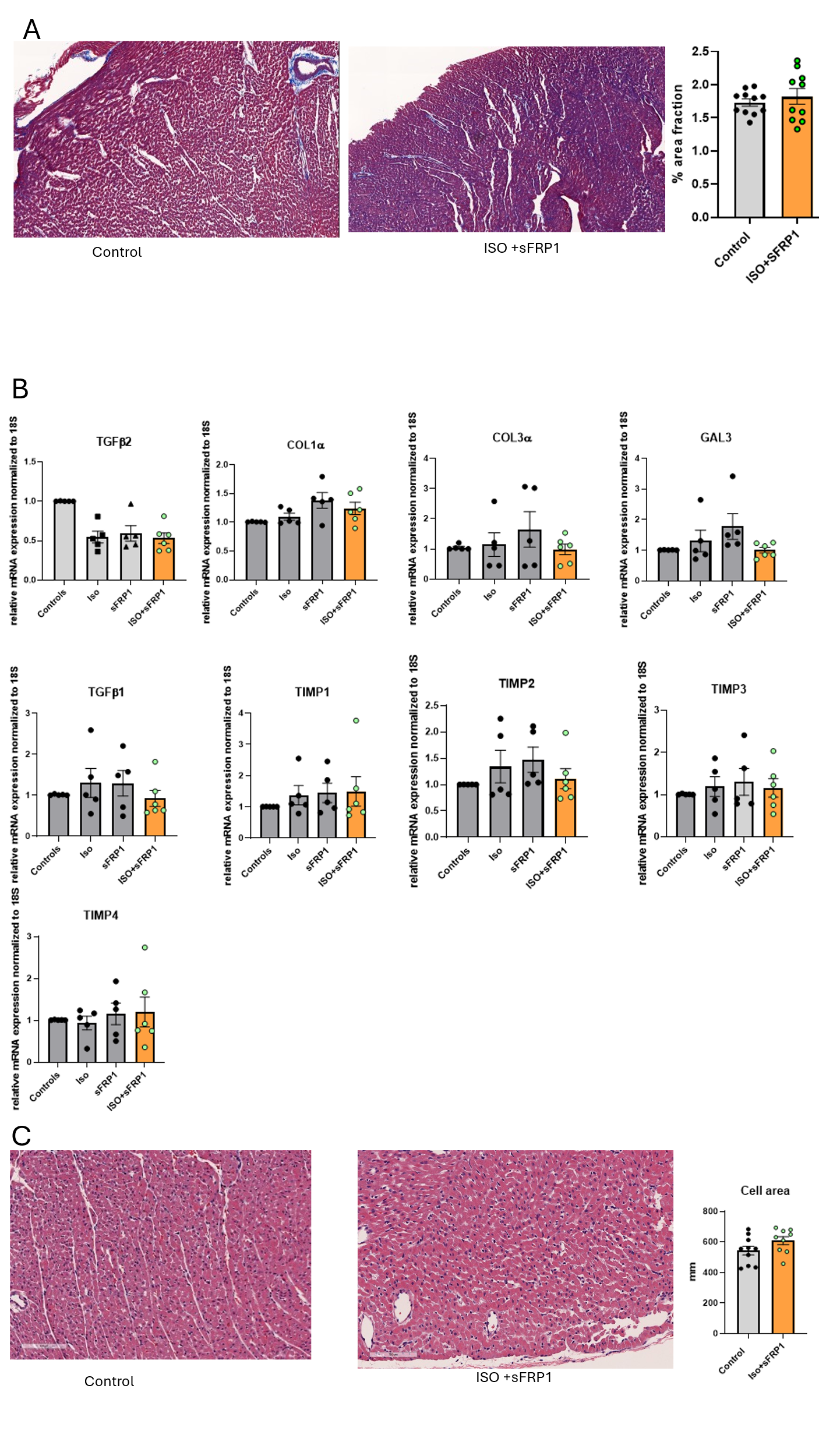

**Supp. Figure 8. Absence of fibrosis and hypertrophy in rats treated with ISO+sFRP1.**

(A) Masson's trichrome staining of vehicle-treated controls and ISO+sFRP1-treated rats. Quantification found on the right of the panel n=10/group.

(B) RT-qPCR of fibrotic gene expression in rats treated with vehicle (Control), ISO, sFRP1 or ISO+sFRP1. Expression of *TGFB2, COL1A1, COL3A1, Galectin-3, TGFB1*, and *TIMP1-4*. Gene expression was normalized to 18S, and data are presented as a relative fold change to Controls. Control n=5, ISO n= 5, sFRP1 n*=5*, ISO+sFRP1 n=6.

(C) Hematoxylin and Eosin staining of vehicle-treated controls, and ISO+sFRP1- treated rats with quantification. n=10/group.

All data are log_2_ transformed and error bar denotes mean± SEM. *P* values are notated in the figure. Fitting a mixed model, Tukey’s multiple comparisons test was used for (B data sets. Unpaired 2-tailed *t*-test was used in (A and C). ISO, Isoproterenol; sFRP1, secreted frizzled protein 1.

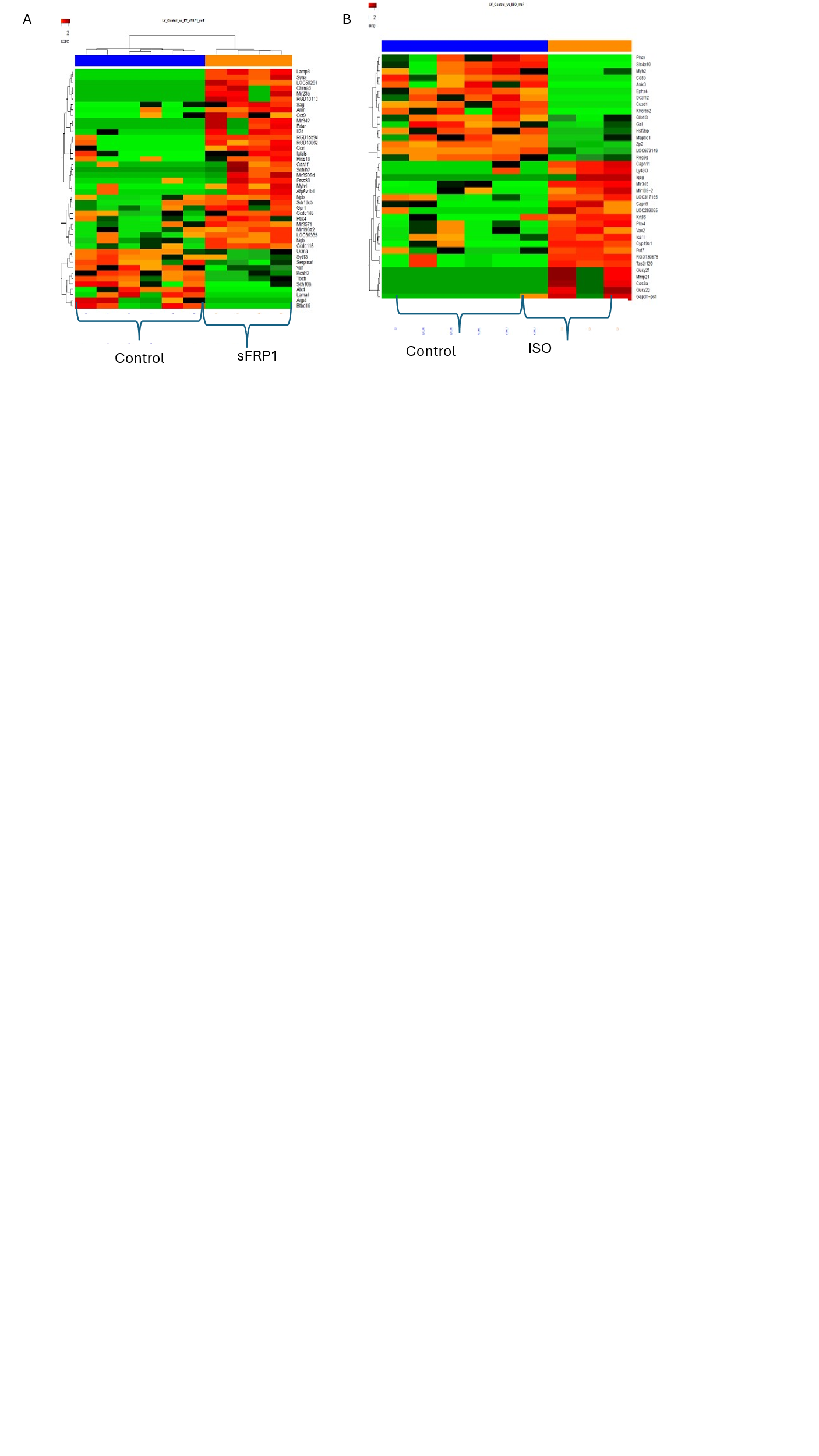

**Supp. Fig. 9 Bulk RNA sequencing analysis of rats treated with ISO or sFRP1.**

(A) Heatmap of the top 50 significantly differentially expressed genes (DEGs) in sFRP1- or (B) ISO-treated rats. Unsupervised hierarchical clustering separated treated from control groups. n=6/group; Wilcoxon rank-sum test was used to compare groups on normalized data.

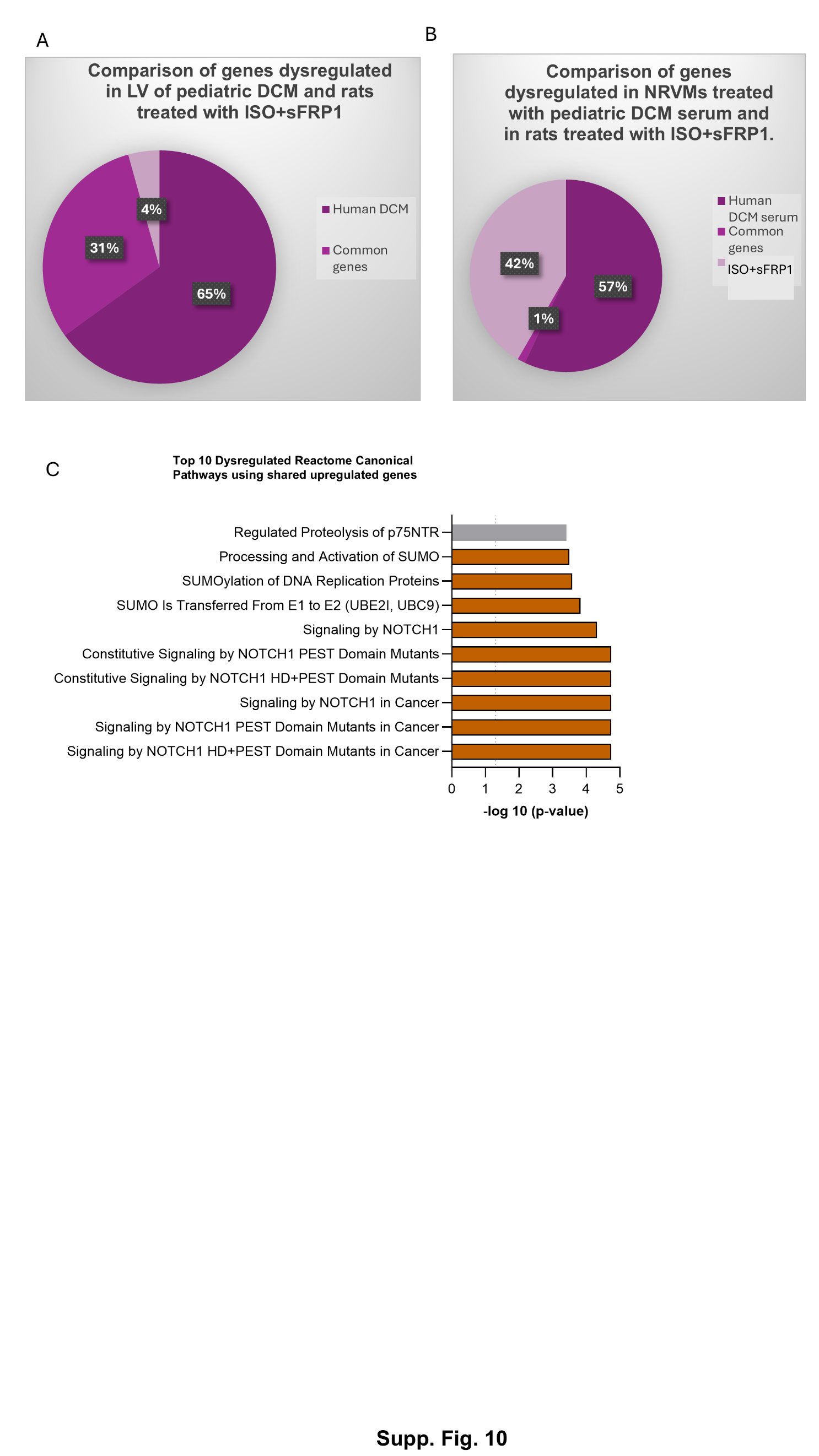

**Supp. Fig. 10. Bulk RNA sequencing analysis of rats treated with ISO+sFRP1 compared to pediatric iDCM hearts and DCM serum treated NRVMs.**

(A) Venn diagram representation of overlapping genes altered in pediatric iDCM left ventricle (LV) tissue compared to non-failing controls, and in LV tissue from ISO+sFRP1-treated rats compared to controls.

(B) Venn diagram representation of overlapping genes altered in pediatric iDCM serum-treated NRVMs compared to cells treated with serum from age-matched non-failing controls (human iDCM serum), and in LV tissue from ISO+sFRP1-treated rats compared to controls.

(C) Pathway analysis using the upregulated genes shared in serum-treated NRVMs and LV tissue of ISO+sFRP1-treated rats identified by Reactome pathway analysis tool.

DCM, dilated cardiomyopathy; ISO-Isoproterenol, sFRP1- secreted frizzled protein 1; NRVMs, neonatal rat ventricular myocytes; LV, left ventricle.

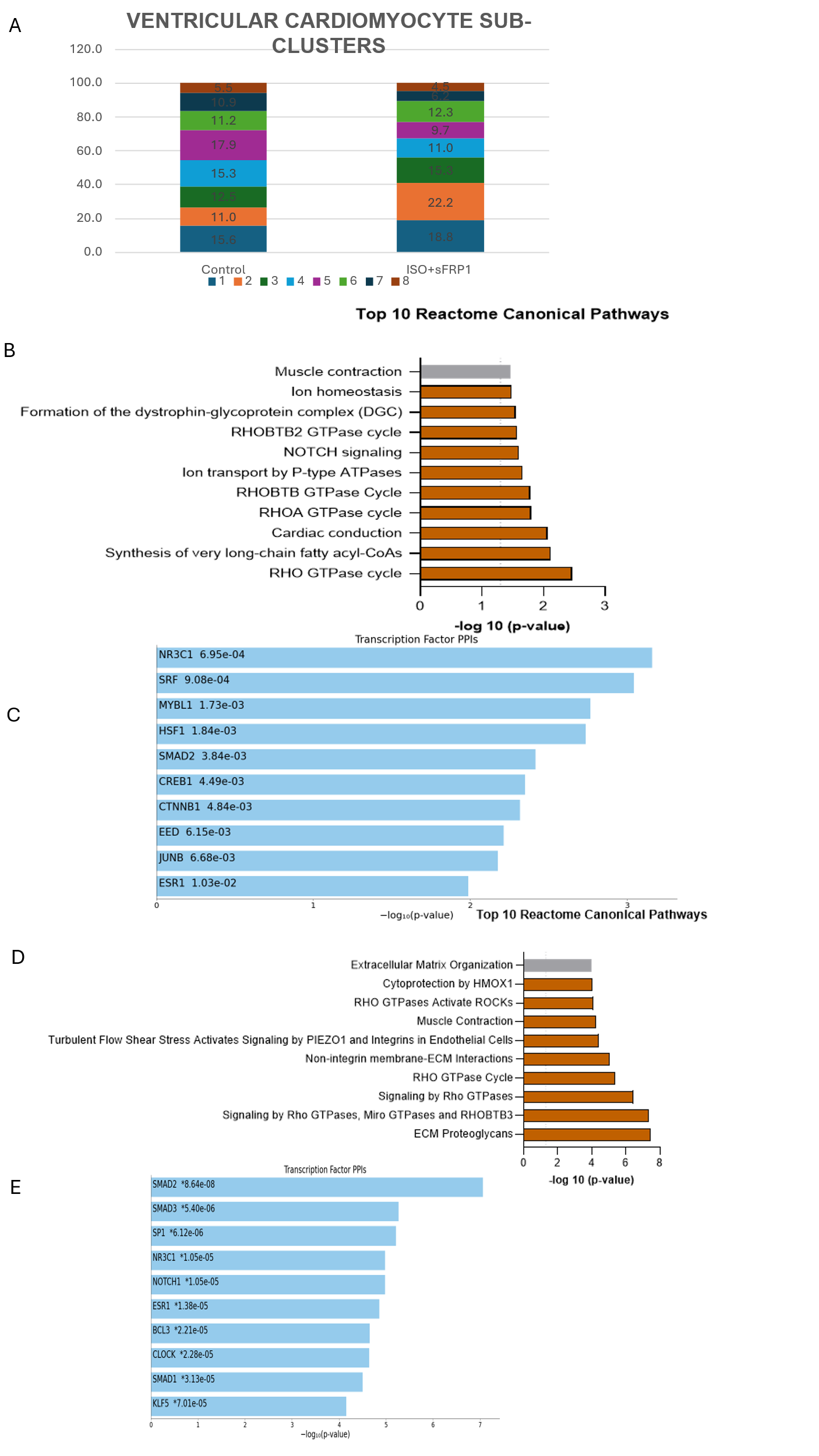

**Supp. Fig. 11. Single nuclei RNA-sequencing analysis of rats treated with ISO+sFRP1 (ventricular cardiomyocyte).**

(A) Ventricular cardiomyocyte sub-clusters from rats treated with ISO+sFRP1 compared to controls.
(B) Reactome pathway analysis using upregulated genes in Ventricular cardiomyocyte sub-cluster 2 of ISO+sFRP1 vs Control rats.
(C) Enrichr analysis identifies transcription factors predicted to control expression of upregulated genes in ventricular cardiomyocyte sub-cluster 2 (right).
(D) Reactome pathways analysis using upregulated genes in ventricular cardiomyocyte sub-cluster 6 of ISO+sFRP1 vs Control rats (left).
(E) Enrichr analysis identifies transcription factors predicted to control expression of upregulated genes in ventricular cardiomyocyte sub-cluster 6 (right).

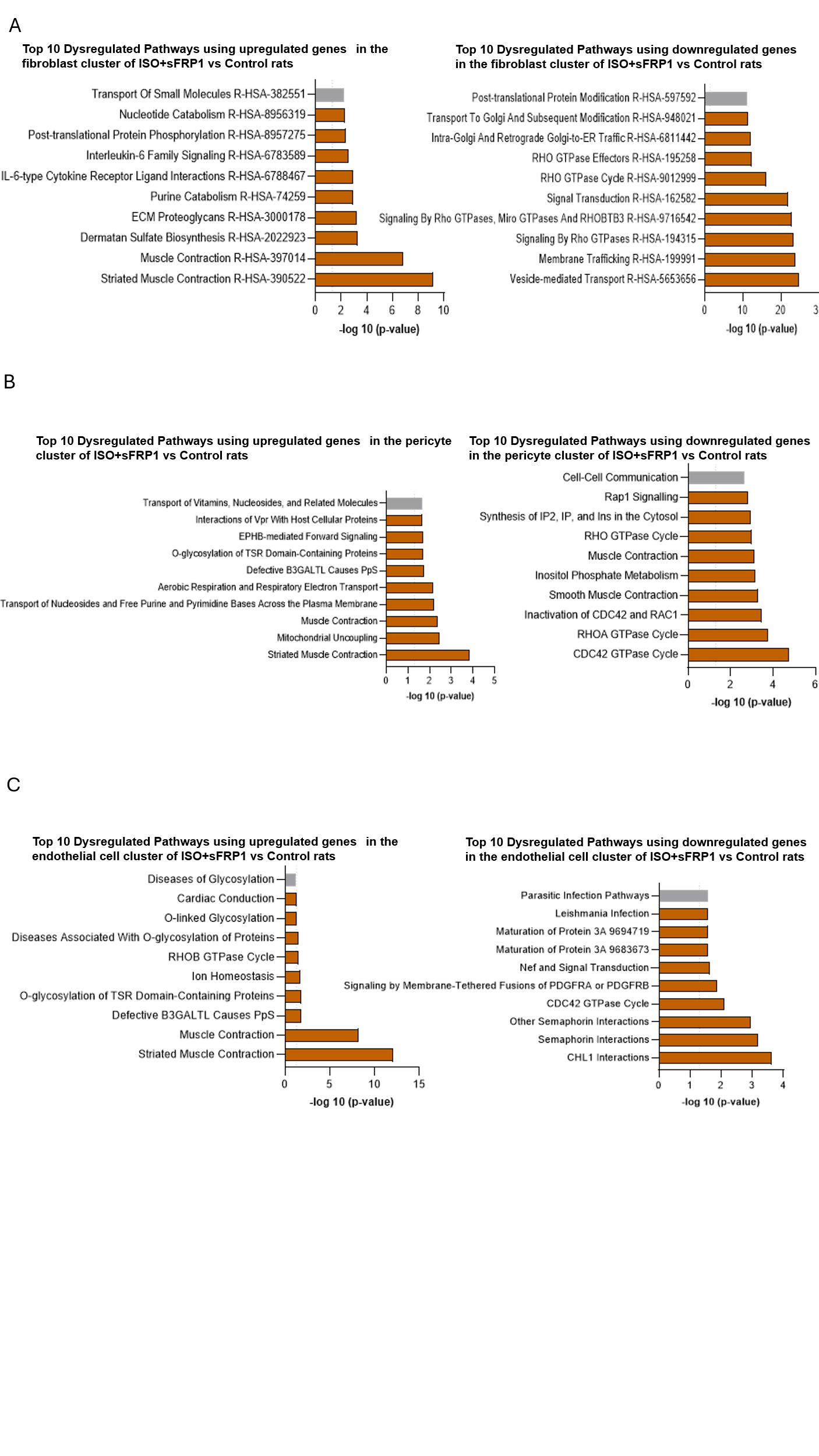

**Supp. Fig. 12. Pathway analysis of differentially expressed genes in single nuclei RNA-sequencing analysis of rats treated with ISO+sFRP1 (stromal cell cluster analysis).**

(A) Pathway analysis using the upregulated (left panel) and downregulated genes (right panel) in the fibroblast cluster of ISO+sFRP1 vs Control rats using Reactome pathways analysis tool

(B) Pathway analysis using the upregulated(left) and downregulated genes(right) in the pericyte cluster of ISO+sFRP1 vs Control rats using Reactome pathways analysis tool

(C) Pathway analysis using the upregulated(left) and downregulated genes(right) in the endothelial cell cluster of ISO+sFRP1 vs Control rats using Reactome pathways analysis tool.

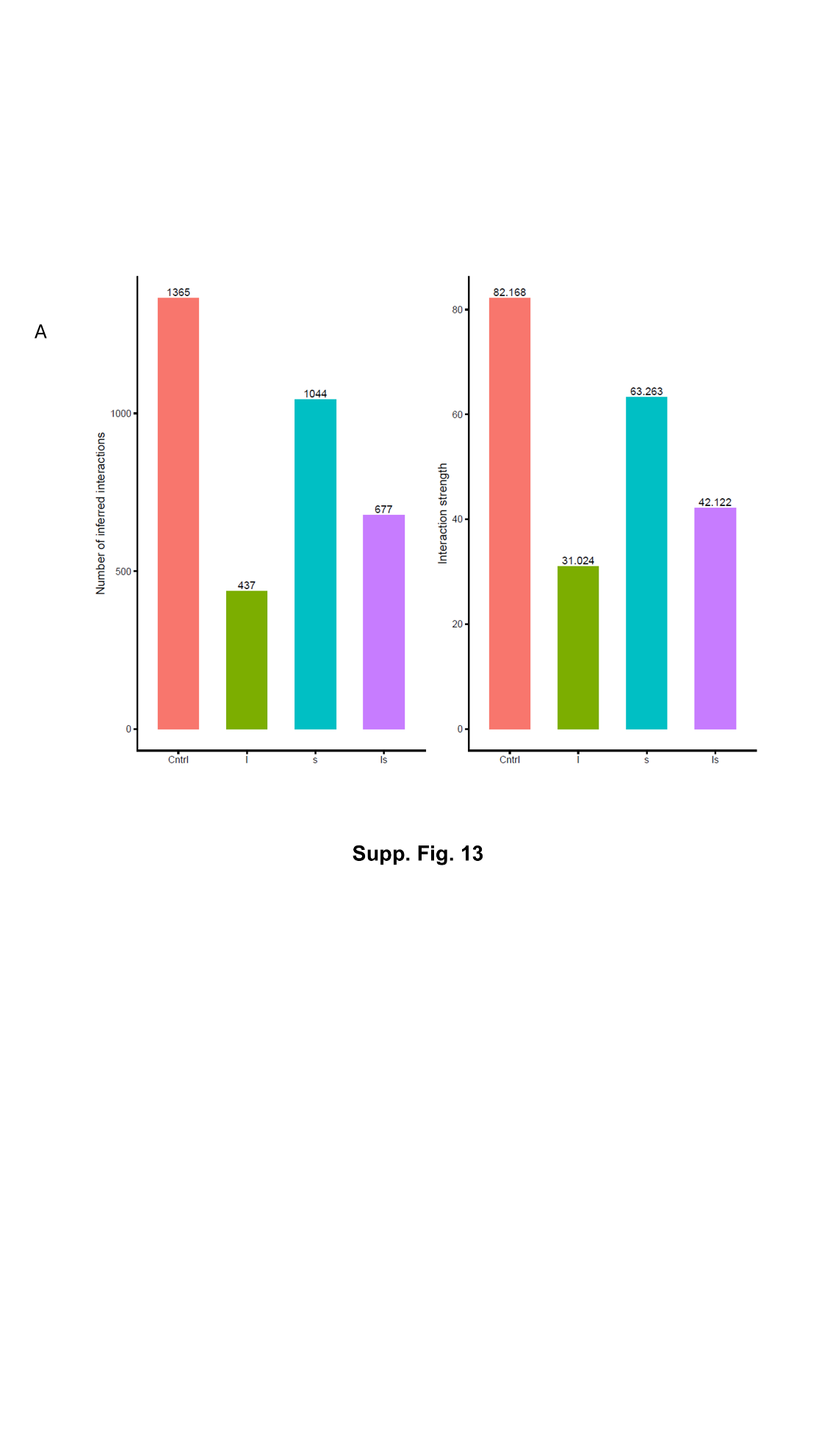

**Supp. Fig.13. Cellchart analysis of intercellular communication in ISO+sFRP1 treated rats compared to vehicle treated controls**

(A) Quantification of global intercellular communication in the ISO+sFRP1 model compared to vehicle-treated controls

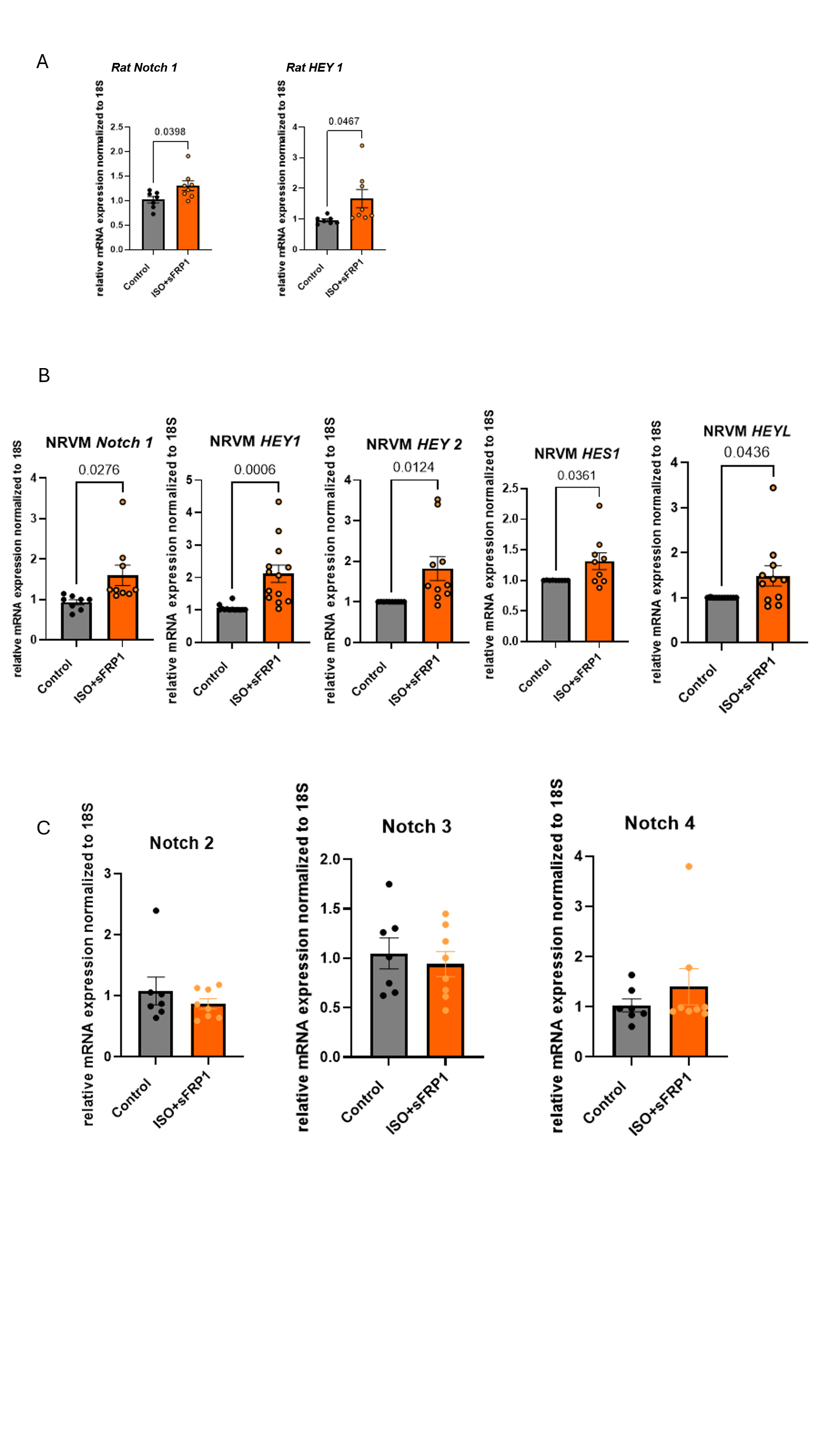

**Supp. Fig.14. ISO+sFRP1-treated Rats and NRVMs have increased Notch 1 gene expression and Notch target genes**

(A) RT-qPCR of Notch1 and Hey1 gene expression in ISO+sFRP1 rat LV tissue compared to Control. Gene expression was normalized to 18S, and data are presented as a relative fold change to Controls. n=8/group.

(B) RT-qPCR of Notch target genes in NRVMs treated with vehicle (Control) and ISO+sFRP1. Expression of *Notch1*, *HEY1, HEY2, HES1* and *HEYL* genes. Gene expression was normalized to 18S, and data are presented as a relative fold change to Controls. n= 6 independent NRVM preps/group.

(C) RT-qPCR of Notch 2-4 expression in NRVMs treated with Control, and ISO+sFRP1 only.

Gene expression was normalized to 18S, and data are presented as a relative fold change to Controls. *n* = 6 independent NRVM preps/group. All data are log_2_ transformed and error bar denotes mean± SEM. Only significant *p* values are notated in the figure. Unpaired t-test was used for all datasets. ISO, Isoproterenol; sFRP1, secreted frizzled protein 1; NRVMs, neonatal rat ventricular myocytes.

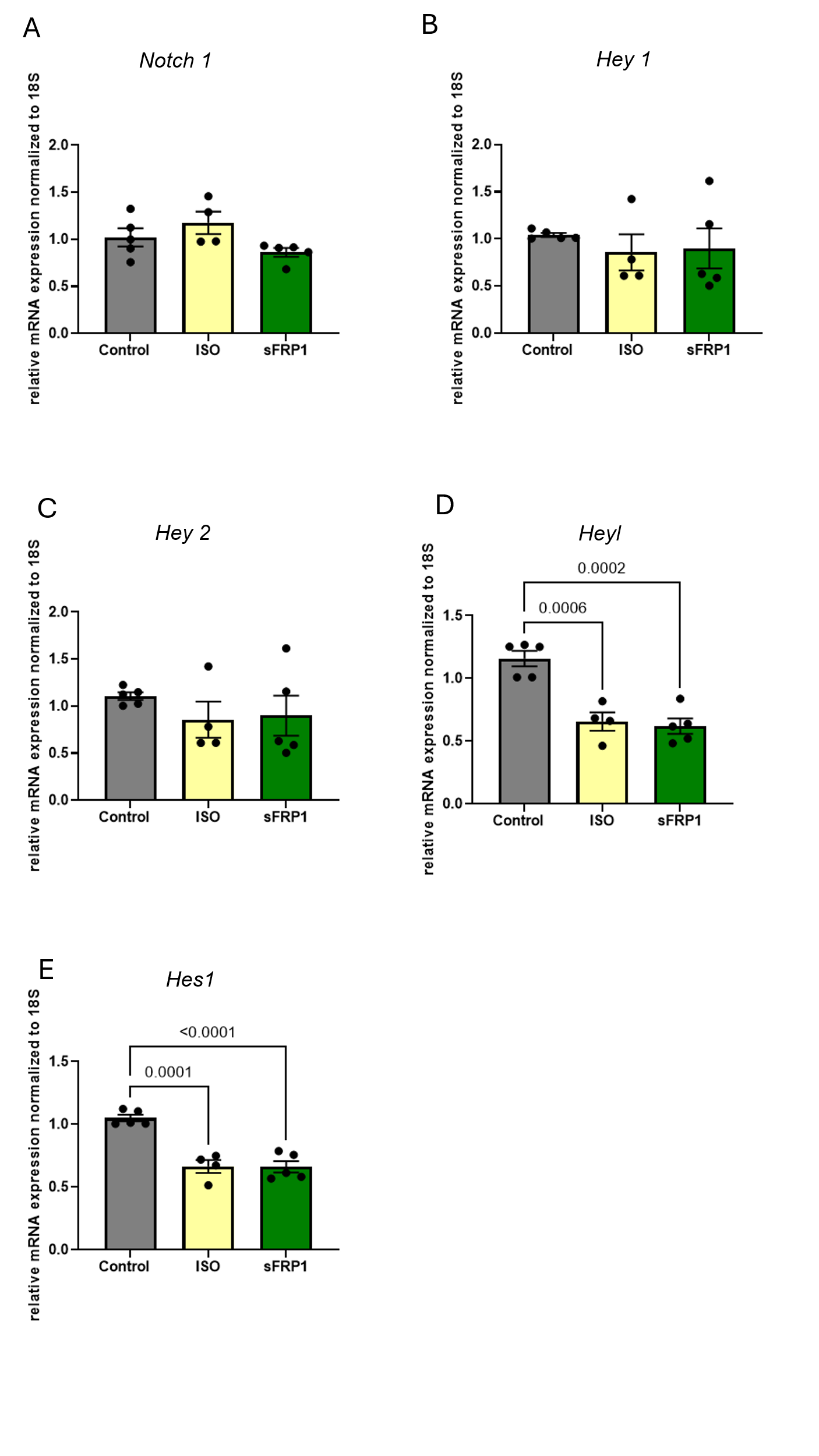

**Supp. Fig.15. ISO or sFRP1-treated NRVMs do not have an increase in expression of Notch genes.**

RT-qPCR of fetal gene program (FGP) expression in NRVMs treated with Control, ISO or sFRP1. Expression of (A) *Notch 1*, (B) *HEY1* (C) *HEY2,* (D) *HEYL* and (E) *HES1*. Gene expression was normalized to 18S, and data are presented as a relative fold change to Controls. n=4 independent NRVM preps/group. All data are log_2_ transformed and error bar denotes mean± SEM. Only significant *p* values are notated in the figure. Fitting a mixed model, Tukey’s multiple comparisons test was used for all data sets. ISO, Isoproterenol; sFRP1, secreted frizzled protein 1; NRVMs, neonatal rat ventricular myocytes.

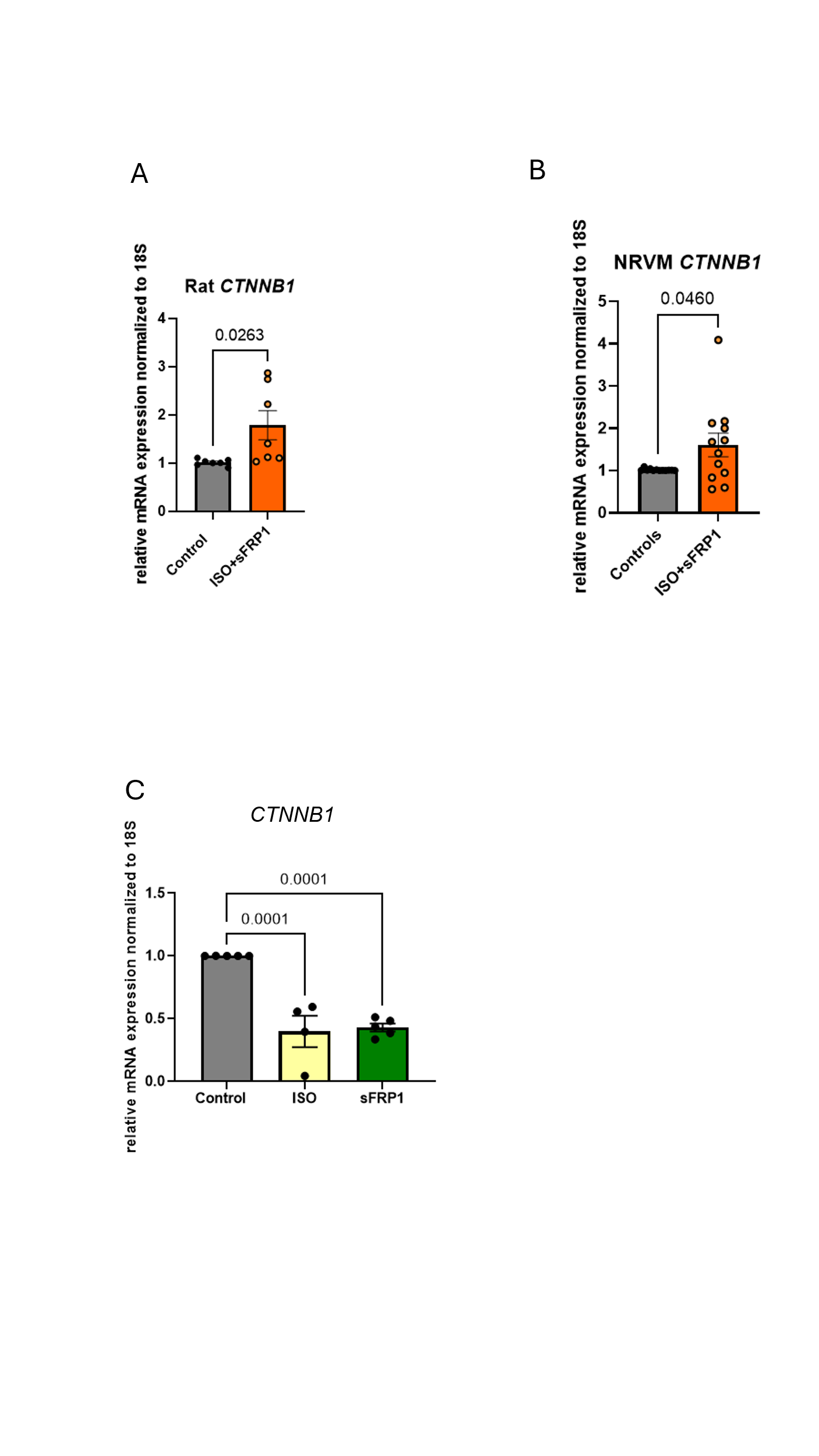

**Supp. Fig.16. ISO+sFRP1-treated Rats and NRVMs have increased *CTNNB1* expression.**

(A) RT-qPCR of CTNNB1 gene expression in ISO+sFRP1-treated rat LV tissue compared to Control. Gene expression was normalized to 18S, and data are presented as a relative fold change to Controls. n=7/group. Unpaired 2-tailed *t*-test was used in analysis.
(B) RT-qPCR of CTNNB1 gene expression in ISO+sFRP1-treated NRVMs to Control. Gene expression was normalized to 18S, and data are presented as a relative fold change to Controls. n=7/group. Unpaired 2-tailed *t*-test was used in analysis.
(C) RT-qPCR of CTNNB1 gene expression in NRVMs treated with vehicle (Control), ISO, or sFRP1. Gene expression was normalized to 18S, and data are presented as a relative fold change to Controls. n=8 independent NRVM preps/group. Fitting a mixed model, Tukey’s multiple comparisons test was in analysis.

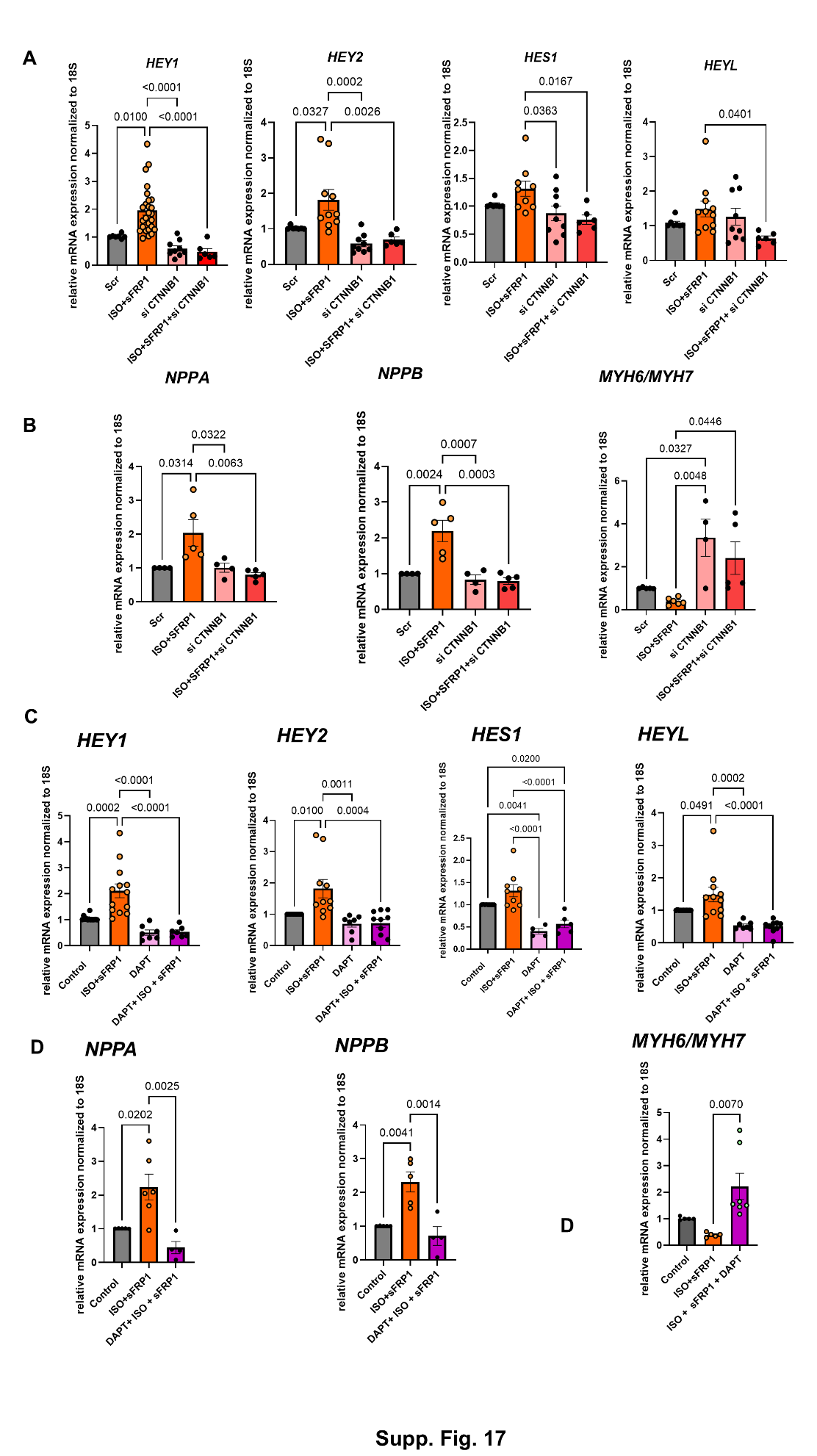

**Supp. Fig.17. Notch and β-catenin inhibition prevent pathological remodeling *in vitro.***

(A) RT-qPCR of Notch target gene expression in NRVMs transfected with scrambled control siRNA (scr) or siCTNNB1, and treated with vehicle (Control) or ISO+sFRP1. Expression of *HEY1, HEY2, HES1* and *HEYL* genes. Gene expression was normalized to 18S, and data are presented as a relative fold change to Controls. n=6 independent NRVM preps/group
(B) RT-qPCR of FGP gene expression in NRVMs transfected with scrambled control siRNA (scr) or siCTNNB1, and treated with vehicle (Control) or ISO+sFRP1. Expression of *NPPA, NPPB,* and ratio of *MYH6* to *MYH7*. Gene expression was normalized to 18S, and data are presented as a relative fold change to Controls. n=4 independent NRVM preps/group
(C) RT-qPCR of Notch target gene expression (*HEY1, HEY2, HES1* and *HEYL)* in NRVMs treated with vehicle (Control), ISO+sFRP1, DAPT (20µM) or ISO+sFRP1+DAPT. Gene expression was normalized to 18S, and data are presented as a relative fold change to Controls. n=7 independent NRVM preps.
(D) RT-qPCR of FGP in NRVMs treated with vehicle (Control), ISO+sFRP1, DAPT (20µM) or ISO+sFRP1+DAPT. Gene expression was normalized to 18S, and data are presented as a relative fold change to Controls. n=5 independent NRVM preps.

Gene expression was normalized to 18S, and data are presented as a relative fold change to Controls. Tukey’s multiple comparisons test was used for all data sets.
